# Mind Captioning: Evolving descriptive text of mental content from human brain activity

**DOI:** 10.1101/2024.04.23.590673

**Authors:** Tomoyasu Horikawa

## Abstract

A central challenge in neuroscience is decoding brain activity to uncover mental content comprising multiple components and their interactions. Despite progress in decoding language-related information from human brain activity, generating comprehensive descriptions of complex mental content associated with structured visual semantics remains challenging. We present a method that generates descriptive text mirroring brain representations via semantic features computed by a deep language model. Constructing linear decoding models to translate brain activity induced by videos into semantic features of corresponding captions, we optimized candidate descriptions by aligning their features with brain-decoded features through word replacement and interpolation. This process yielded well-structured descriptions faithfully capturing viewed content, even without relying on the canonical language network, thereby revealing explicit representations of fine-grained structured semantic information outside this network. The method also successfully generalized to verbalize recalled content, demonstrating the potential for non-verbal thought-based brain-to-text communication, which could aid individuals with language expression difficulties.

## Introduction

Humans can recognize and recall intricate visual content comprising multiple semantic components, including objects, places, actions, and events, along with their interactions and relationships. These elaborate and structured mental representations form the foundation for translating thoughts into language and communicating experiences with others. Recently, substantial progress has been made in brain decoding of language-related information, enabling the direct production of linguistic outputs, such as text, from the human brain^1–4^. However, decoding the perceptual—and not to mention mental—content associated with visual semantics to generate comprehensive descriptions of subjective experiences remains challenging. Translating brain activity linked to non-linguistic semantic information, or thoughts, into verbal descriptions could significantly enhance our ability to interpret diverse mental states, opening up numerous possibilities for applications, particularly with text-prompt-based systems (e.g., ChatGPT^5^ and Gemini^6^), as well as for scientific research.

Prior research on decoding visual semantics using human functional magnetic resonance imaging (fMRI) has focused on individual components or static images. This narrow focus has hindered the decoding of complex content involving interactions between multiple elements, thus obscuring our understanding of how the brain represents rich and structured visual semantics. While studies have successfully decoded individual components in viewed^7–9^, imagined^7^, and dreamed^10^ content using object-or word-level features, they have fallen short of capturing neural representations of interactions and relationships that are not predicted by their individual components^11^ and are crucial for recognizing actions and social interactions^12–16^.

Some researchers have incorporated caption databases^17^ or deep neural network (DNN)-based modules, such as non-linear image captioning models^18–20^, to produce sentence-level decoding predictions that appear to have linguistic structure. However, predictions based on database search methods are limited to existing, often deliberately structured, descriptions that may not capture the full complexity of diverse visual content. In addition, non-linear methods can introduce spurious information not “explicitly represented” in the brain^21–23^. Specifically, non-linear captioning models can construct sentence-like structured outputs even from object-level visual features^24^, which inherently lack relational information Therefore, these approaches, designed to produce linguistically structured outputs, are not ideal for examining whether structured visual semantics, essential for representing relational information, are genuinely encoded in the brain through decoder outputs.

To overcome these limitations, we introduce a new generative decoding method called *mind captioning*, which generates descriptive text mirroring semantic information represented in the brain (Fig. 1). Our method combines linear feature decoding analysis^7,9,10^, using semantic features computed by a deep language model (LM), with a novel optimization method that generates text based on these features. Semantic features serve as intermediate representations for decoding (translating) semantic information from the brain into text. They can act as a bridge for decoding both perceptual and mental content, as shared representations exist between visual perception and mental imagery, particularly for high-level information^7,10,25–27^. Additionally, deep LMs offer the advantage of effectively capturing contextual meanings, which are crucial for delineating intricate interrelationships^28–33^.

**Fig. 1.**
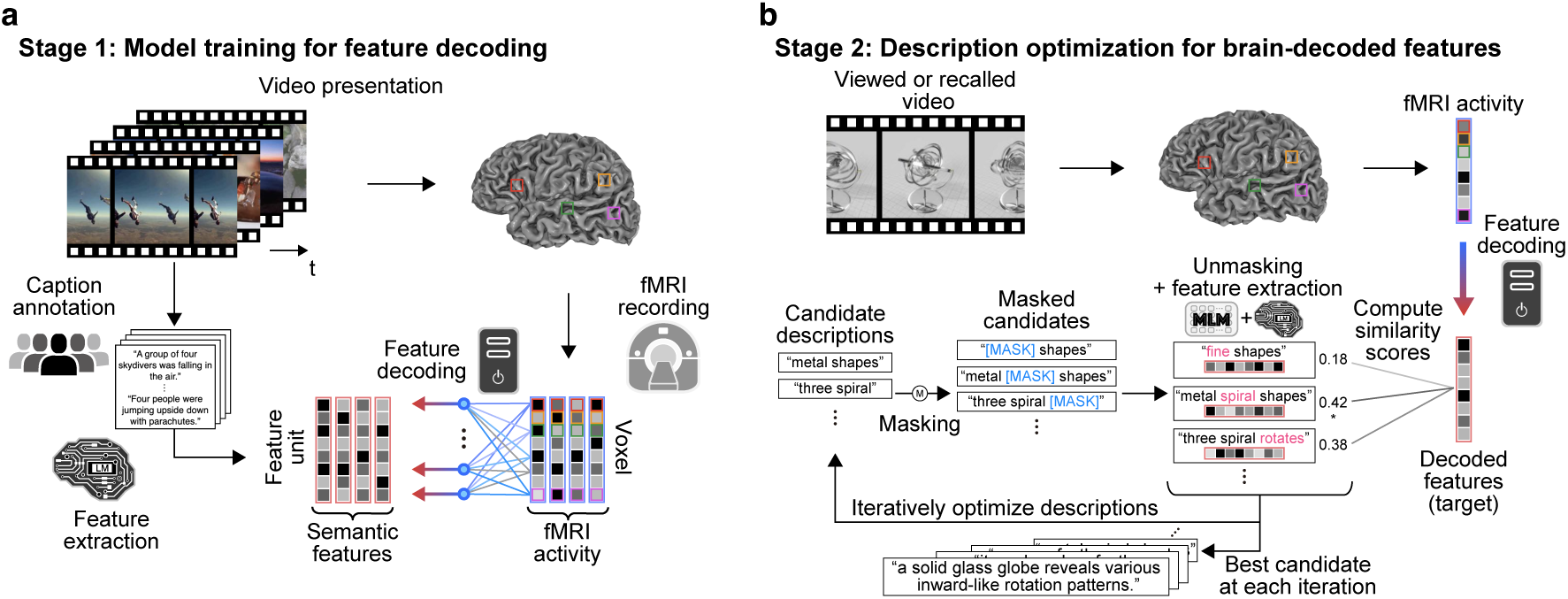
Mind captioning. Our method consisted of two stages. **a**, We first trained linear decoding models to decode whole-brain fMRI activity, measured while each subject viewed videos, into semantic features from the captions of the videos using an LM (frozen). **b**, We then used those models to decode brain activity induced by novel video stimuli or by recall-based mental imagery of those videos, and optimized candidate descriptions iteratively by aligning their features with brain-decoded features through word replacement and interpolation, leveraging another LM pre-trained for MLM (frozen). The optimization consisted of three stages: masking, unmasking, and candidate selection. During masking, we randomly applied masks by replacing words with a mask or interpolating masks into the candidate word sequences. During unmasking, the MLM model created new candidates by filling in the masks in the masked candidates based on the context of surrounding words. During candidate selection, we computed semantic features of all new and original candidates using the LM for feature extraction. We then evaluated the similarity between those candidate features and the target brain-decoded features to select the top candidates for further optimization. The optimization process was initiated from a non-informative word (“ <UNK>”) to avoid incorporating any prior assumptions for description generation and was repeated 100 times. See Extended Data Fig. 2 for details on the model and parameter validations.

The challenge lies in linguistically interpreting the information in semantic features decoded from brain activity. Although the ideal approach would involve examining all possible word sequences to identify the description whose semantic features best match the decoded features, this is not feasible because the number of candidates is infinite.

We thus developed an iterative optimization method that generates descriptive text from scratch by progressively aligning the semantic features of candidate descriptions with target brain-decoded features through word replacement and interpolation in a search for the best description (Fig. 1b). Crucially, we leveraged an LM pre-trained for masked language modeling (MLM)^34^ to constrain the search space during optimization. By directly optimizing word sequences to match brain-decoded features, our method minimizes dependence on external resources such as caption databases or non-linear captioning models, thereby ensuring the generation of descriptions more closely aligned with brain representations while maintaining the interpretability of structured visual semantics in the brain.

To demonstrate the effectiveness of our method, we first validated it for perceptual content by constructing decoding models (decoders) from stimulus-induced brain activity and then tested their generalizability to activity during recall-based mental imagery. Specifically, we measured brain activity in six subjects using fMRI while they viewed or recalled video clips^35^ (Extended Data Fig. 1) and created a data sample by averaging fMRI volumes measured during viewing and recalling each video. To enhance the quality of the fMRI data, we averaged data samples over five repetitions for each stimulus or imagery item in the test phase. Using the samples from stimulus-induced brain activity, we trained decoders to predict semantic features, which were computed from corresponding video captions using an LM (DeBERTa-large^36^). We then used these decoders to translate brain activity associated with viewed and recalled content into semantic features for novel test videos not used during training. Finally, we used the decoded features to optimize the text by employing an MLM model (RoBERTa-large^37^). Through these analyses, we aimed to validate our method’s capability to generate comprehensive descriptions of both viewed and recalled content from brain activity, establishing a novel framework for decoding non-verbal mental content and exploring the neural basis of structured visual semantics.

## Results

### Generating viewed content descriptions

The optimization of text, based on decoded features from stimulus-induced brain activity, resulted in a progressive evolution of descriptive texts (Fig. 2a). Initially, the descriptions were fragmented and lacked clear meaning. However, through iterative optimization, these descriptions naturally evolved to have a coherent structure and effectively capture the key aspects of the viewed videos. Notably, the resultant descriptions accurately reflected the content, including the dynamic changes in the viewed events (Fig. 2b). Furthermore, even when specific objects were not correctly identified, the descriptions still successfully conveyed the presence of interactions among multiple objects (e.g., Fig. 2b bottom left).

**Fig. 2.**
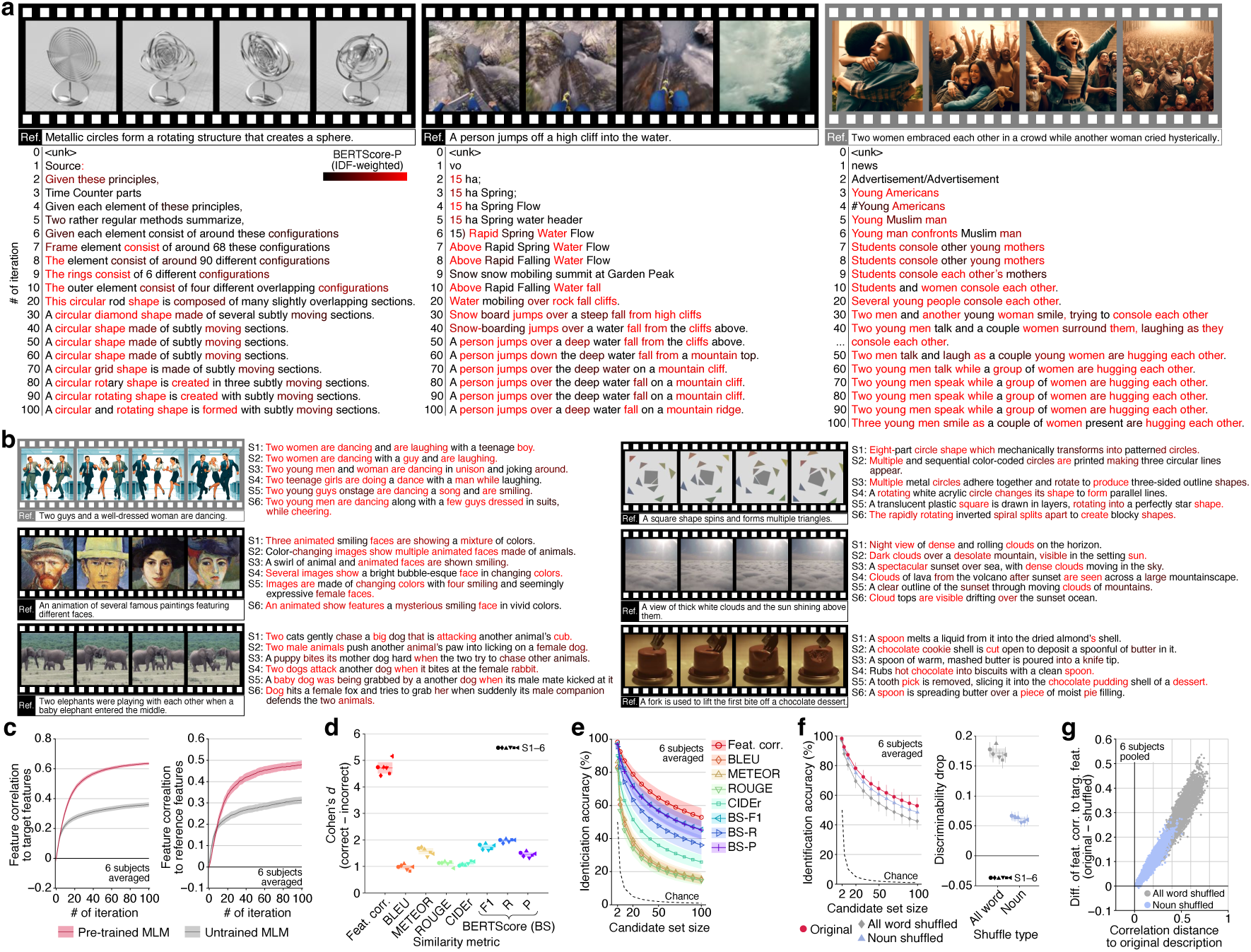
Generating viewed content descriptions. Descriptions were generated using features from all LM layers decoded from whole-brain activity. **a**, Evolved descriptions during the optimization (see Supplementary Video 1 for more example). **b**, Descriptions after 100 iterations for all subjects (see Extended Data Fig. 3a and Supplementary Video 2 for more example). In (**a**), (**b**), the color indicates accuracy [inverse document frequency (IDF)-weighted BERTScore-P]. A reference caption of the video is shown below frames. **c**, Feature correlations between features of generated descriptions and those decoded from the brain, as well as those computed from correct references. **d**, Cohen’s *d* of discriminability (see Extended Data Fig. 4b for raw scores). **e**, Video identification accuracy with varying numbers of candidates. **f**, Effects of word-order shuffling on video identification accuracy and discriminability. **g**, Scatterplot of the correlation distances (one minus feature correlation) between the original and shuffled descriptions against the difference in feature correlations to target features between original and shuffled descriptions. Each dot indicates a shuffled description. Shades in (**c**), (**e**), and error bars in (**d**), (**f**) indicate 95% confidence intervals (C.I.) across samples (*n* = 72). Shades in (**d**), (**f**) indicate 95% C.I. across subjects (*n* = 6). See Extended Data Fig. 4 for individual results. [Note: To comply with bioRxiv’s policy on displaying human faces, frames containing real human faces in this figure (framed by gray) have been replaced with images synthesized by DALL·E3 (https://openai.com/dall-e-3), based on captions annotated by humans for the video.]

Throughout the optimization process, the features of the evolved descriptions exhibited increasingly stronger correlations with the target features and, consequently, with the features of the reference captions annotated on the viewed videos (Fig. 2c). In contrast, the same optimization performed with an untrained MLM model, which randomly suggests candidate words during unmasking—a process analogous to a genetic algorithm—did not yield comparable results (Extended Data Fig. 3b). These results emphasize the significance of contextual information from a pre-trained MLM model to efficiently explore descriptions closely aligned with brain representations and to enhance description quality.

To assess decoding performance, we used multiple similarity metrics to evaluate the similarity between the generated descriptions and the reference captions. We calculated similarity scores for both the captions of the viewed (correct) and irrelevant (incorrect) videos and defined discriminability as the difference between them. The generated descriptions exhibited significantly high discriminability across all metrics and subjects (Fig. 2d; Wilcoxon signed-rank test, one-tailed, *P* < 0.01, FDR corrected across metrics and subjects), indicating that these descriptions were accurate enough to differentiate video content.

To gain an intuitive understanding of the performance, we evaluated the accuracy of video identification by comparing the generated descriptions to both correct and incorrect captions across various numbers of candidates. Performance consistently exceeded chance levels for all set sizes, with approximately 50% accuracy among 100 distinct possibilities for all subjects when using feature correlation (chance = 1%; Fig. 2e), demonstrating the effectiveness of our method in translating detailed information from the brain into text through semantic features.

To further highlight the effectiveness of our method, it is worth noting that it surpasses the previous approach and holds promise for continued improvement. Our method outperformed the database search method^17^ in capturing subjective experience more accurately and flexibly (Extended Data Fig. 5), as its generated descriptions more closely aligned with captions that individual subjects rated as highly consistent with their own perceptions (Extended Data Fig. 1a and 5e). Furthermore, we showed that our method robustly generated descriptions that faithfully reflected the viewed content, regardless of the LMs employed, and found correlations between brain encoding performance—a metric assessing alignment between the brain and models^31,38^—and the text generation performance of the LMs (Extended Data Fig. 6). These results suggest that employing LMs more closely aligned with the brain (e.g., GPT3^5^, OPT^39^, and LLaMA^40^) may further improve the effectiveness of our method.

### Capturing visual relational information

A key advantage of structured descriptions over simple word lists is their ability to organize words to convey contextual meaning, including relational information between components (e.g., the differences between “a bird eats a snake” and “a snake eats a bird”)^41^. To examine if our generated descriptions accurately captured such visual relational information through proper word arrangement, we assessed the effect of shuffling their word order—either for all words or nouns only (up to 1,000 shuffled variants; see Methods)—under the hypothesis that if the original word order accurately conveyed visual relationships, shuffling would reduce discriminability.

While the shuffled descriptions retained reasonably high accuracy in identifying videos—indicating that word lists alone provide informative cues—shuffling all words, or even just the nouns, significantly impaired discriminability (Fig. 2f; Wilcoxon signed-rank test, one-tailed, *P* < 0.01, FDR corrected across subjects). This reduction in discriminability remained robust even when using the minimally disrupted shuffled descriptions, which had the highest fluency (or linguistic acceptability) as assessed by the pseudo-log-likelihood score from MLM scoring^42,43^ (Extended Data Fig. 4f). These results demonstrate that our method generates descriptions that capture more detailed information than simple word lists.

Interestingly, the impact of shuffling was more pronounced when using features from deeper LM layers to generate descriptions (Extended Data Fig. 7), underscoring the importance of the deep structure of the LMs in constructing contextual semantic representations and accurately decoding relational information.

### Validating structured semantics decoded from the brain

To validate that our method genuinely exploited the structured visual semantics encoded in the brain, it is crucial to ensure that the coherent structure of the generated descriptions—key to depicting visual relations—was not artificially imposed by the MLM model used in the text optimization process (Fig. 1b). We reasoned that if a generated description truly reflects the brain representation of specific structured semantic information—uniquely conveyed by its particular word order rather than by alternative arrangements of the same words—then the target brain-decoded features should exhibit greater similarity to the features of the generated description than to those of shuffled variants. We tested this by comparing feature correlations between the brain-decoded features and both the original and shuffled descriptions.

Supporting our reasoning, shuffling the word order significantly reduced feature correlations, especially when it substantially altered the original meanings (Fig. 2g). The original descriptions scored highest among all variants generated through all-word shuffling and ranked in the top 0.001% for noun-only shuffled variants (six subjects pooled). These results suggest that the generated word sequences authentically reflect the semantic information encoded in the brain.

To further rule out the possibility that the MLM model imposed structured semantics on the generated descriptions, we examined the outputs of decoders trained with semantic features from word-order-shuffled captions—a method intended to prevent the decoders from learning structured semantics. As a result, we confirmed that incoherent word sequences were generated, though they still contained words that semantically matched individual components of the viewed videos (Extended Data Fig. 3c).

Together, these results reinforce our finding that the text optimization process does not impose restrictive constraints on forming linguistically structured outputs when generating coherent and structured descriptions from brain activity. Instead, the observed coherence likely reflects the structured semantics inherently encoded in the brain-decoded features rather than being artificially imposed by the MLM model.

### Contributions from different brain areas

Having confirmed the ability to generate accurate and well-structured descriptions from whole-brain activity, we next examined the contributions of specific brain regions to this decoding, focusing on whether complex structured semantics can be derived independently of the language network. Although numerous studies have shown that a broadly distributed semantic network encodes the meanings of both visual and linguistic information^44–47^, most have focused on category-or word-level representations. Research on structured semantics has primarily concentrated on language processing, often associated with the fronto-temporal language network^48–50^, whereas the neural substrates for structured visual semantics remain comparatively underexplored. Emerging evidence suggests that, while the language network is also recruited in processing meaningful visual scenes^51,52^, other regions—particularly the lateral occipital temporal cortex—encode certain relational aspects of visual information, including interactions and the directedness of actions among persons and objects^12–16^. These findings suggest the intriguing potential to enable the direct decoding and communication of rich structured semantic information from brain activity, bypassing the linguistic processing typically required to translate thoughts into words. Pursuing this possibility could broaden the scope of brain–machine interfaces (BMIs) for converting non-verbal thoughts into text, opening new avenues for semantic decoding that do not rely on language.

To establish a foundation for evaluating decoding performance, we began by analyzing how the brain encodes structured visual semantic information in videos. We constructed two encoding models: one based on the semantic features used in our decoding analysis and the other on visual features from a video recognition DNN pre-trained to classify object and action categories^53^. The semantic encoding model effectively predicted brain activity in the language network and in regions involved in recognizing objects, actions, and interactions^11–16^, spanning the higher visual cortex (HVC) and extended areas of the parietal cortex (Fig. 3a). While the visual model performed better in the lower visual cortex (LVC, V1–V3), the semantic model progressively outperformed it in the HVC and language network (Fig. 3b–e). The shift in relative superiority between the visual and semantic models occurs at the midpoint of the category-selective regions, between their posterior and anterior halves, suggesting a functional boundary (Fig. 3b).

**Fig. 3.**
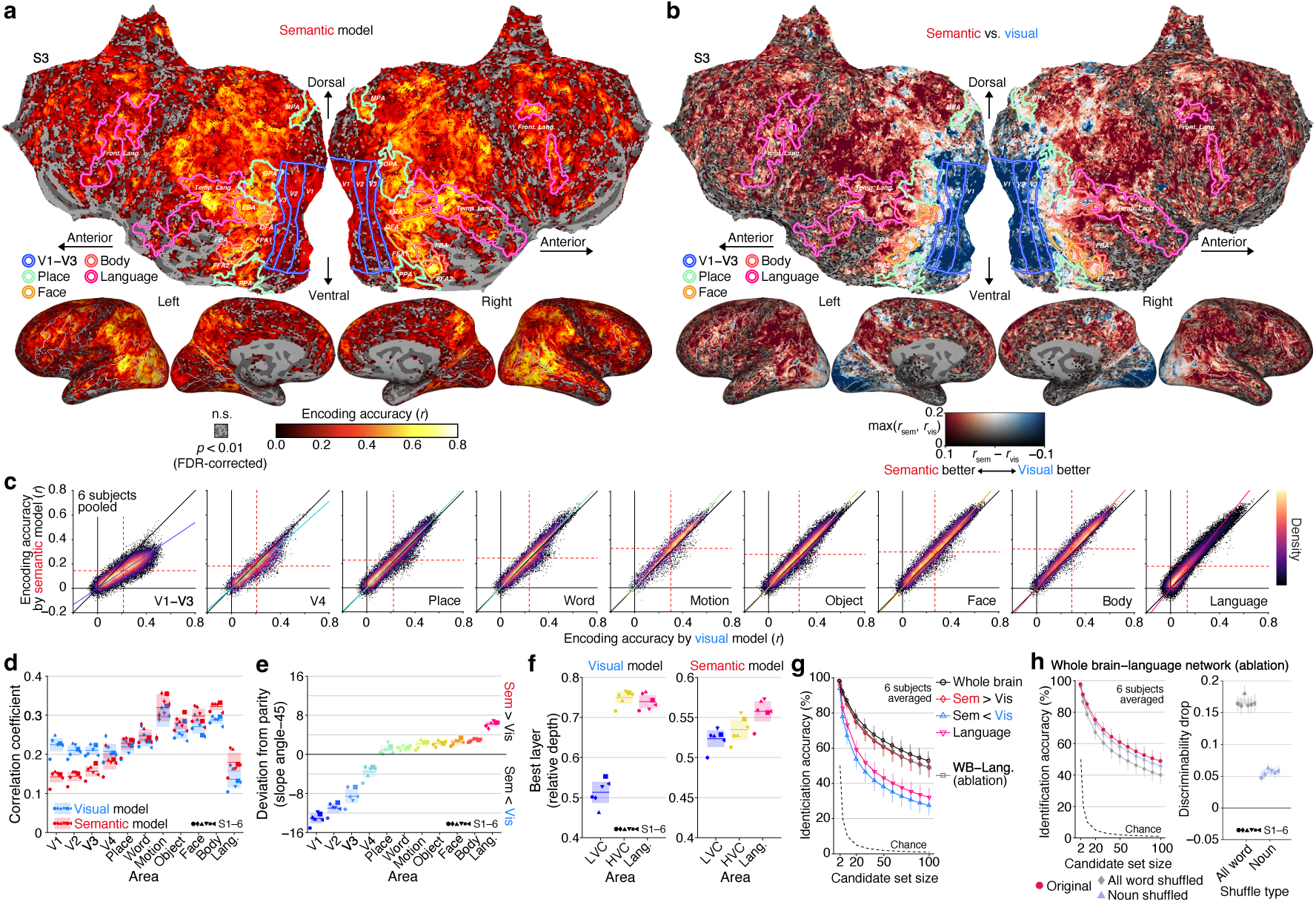
Contributions from different brain areas. A cross-validation analysis was performed within training perception data. Encoding models were trained using features from each layer to generate predictions from multiple layers. The final predictions were constructed based on the best layer per voxel determined using nested cross-validation. **a**, Encoding performance of the semantic model. **b**, Performance difference between the semantic and visual models. **c**, Density heatmap of accuracy (solid lines, best linear fit). **d**, Mean encoding accuracy within each area. **e**, Deviation from parity based on the slope angles of the best linear fit. **f**, Mean of the best layers. The indices of layers with the highest performance were averaged across voxels in each area and then converted into relative depth. **g**, Video identification accuracy from different brain areas. Decoders were trained using voxels with higher encoding accuracy according to the semantic or visual models, voxels within the language network, or voxels from the entire brain except for the language network. **h**, Effects of word-order shuffling on video identification accuracy and discriminability without using the language network. In (**d**)–(**h**), error bars and shades indicate 95% C.I. across samples and subjects, respectively. See Extended Data Fig. 4 and 8 for individual results.

Additionally, voxels in the HVC and language network were better predicted by features from deeper layers of the visual DNN and the LM (Fig. 3f). Notably, the LM layers yielding the highest performance for language network voxels were much deeper than those for the HVC (Wilcoxon rank-sum test, one-tailed, *P* < 0.01, FDR corrected across subjects), highlighting a significant link between the language network and contextualized semantic information. These results demonstrate that the language network, along with other regions, is involved in encoding contextual semantics, consistent with previous studies on its activation by non-verbal visual semantics^51,52^.

We then assessed decoding performance using video identification analysis based on descriptions generated from these brain regions with varying selectivity (Fig. 3g). Focusing on voxels better predicted by the semantic model yielded higher performance, approaching whole-brain activity results for some subjects (Extended Data Fig. 4h). In contrast, decoding from voxels better predicted by the visual model resulted in weaker performance, indicating limited contributions from these voxels even though they are widely distributed across the posterior side of visual category-selective areas. Using only the language network, despite its involvement in encoding contextual semantics, did not produce high performance, suggesting that its contribution may be more supportive than essential. Notably, ablating the language network did not profoundly impact performance, achieving almost 50% accuracy from 100 possibilities (chance = 1%; six subjects averaged), showing decreased accuracy and discriminability by word order shuffling (Fig. 3h; Wilcoxon signed-rank test, one-tailed, *P* < 0.01, FDR corrected across subjects), and even generating intelligible descriptions (Extended Data Fig. 3d).

These results suggest that accurate descriptions capturing structured semantics can be generated without relying on the language network, indicating that structured visual semantic information is represented across regions extending from the anterior portions of the occipital visual cortex to areas outside the language network. Such representations may underlie the comprehension of complex visual events in individuals with global aphasia^51^. These findings also provide further support for the distinction between language and non-verbal thought^54^.

### Generating recalled content descriptions

Finally, we investigated whether the decoders trained on brain activity induced by visual stimuli could be used to generate descriptions of mental content based on brain activity induced by mental imagery of recalled videos, applying the same evaluation procedures as in the perception data analysis. The analysis successfully generated descriptions that accurately reflected the content of the recalled videos, although accuracy varied among individuals (Fig. 4a). These descriptions were more similar to the captions of the recalled videos than to irrelevant ones (Extended Data Fig. 9a and b), with proficient subjects achieving nearly 40% accuracy in identifying recalled videos from 100 candidates (Fig. 4b; chance = 1%). Shuffling word order resulted in a notable reduction in video identification accuracy and discriminability (Fig. 4c; Wilcoxon signed-rank test, one-tailed, *P* < 0.01, FDR corrected across subjects). Excluding the language network from the analysis slightly, but not substantially, reduced video identification accuracy (Fig. 4d).

**Fig. 4.**
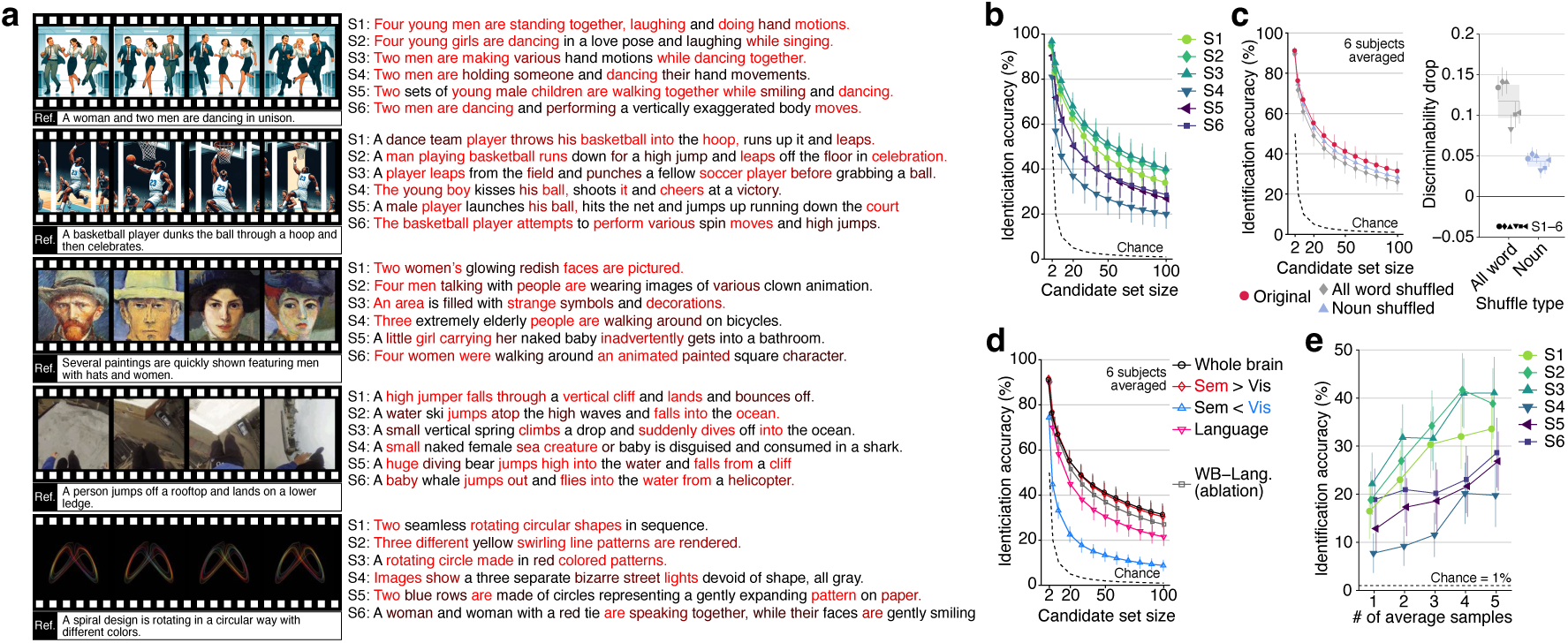
Generating recalled content descriptions. We used the decoders trained on stimulus-induced brain activity (DeBERTa-large; all layers; decoded from whole-brain activity, unless otherwise stated) to analyze the brain activity of subjects engaged in recalling video scenes from memory, prompted by verbal descriptions (see Extended Data Fig. 1b). **a**, Descriptions of recalled content generated after 100 iterations for all subjects (see Extended Data Fig. 3e and Supplementary Video 3 for more examples). **b**, Identification accuracy of recalled videos from individual subjects. **c**, Effects of word-order shuffling on identification accuracy and discriminability of recalled videos. **d**, Identification accuracy of recalled videos from different brain areas. **e**, Identification accuracy of recalled videos with a varying number of averaged samples. For (**a**), (**c**), and (**d**), conventions are the same as for Fig. 2b, f, and Fig. 3g. See Extended Data Fig. 9 and 10 for more data. [Note: To comply with bioRxiv’s policy on displaying human faces, frames containing real human faces in this figure (framed by gray) have been replaced with images synthesized by DALL·E3 (https://openai.com/dall-e-3), based on captions annotated by humans for the video.]

Notably, when the same text generation analysis was applied to brain activity during the preparation period (Extended Data Fig. 1b), all subjects demonstrated lower accuracy compared to the imagery period, with most subjects (except S2) performing at levels closer to chance (Extended Data Fig. 9d). This finding suggests that volitional mental imagery, rather than text reading of verbal cues, is essential for eliciting semantic neural representations, enabling the accurate generation of target content descriptions.

Collectively, these results confirm the generalizability of the decoders to generate descriptions of recalled content, showcasing their capability to effectively verbalize mental representations. Additionally, we were able to obtain comprehensible descriptions of the recalled content with reasonable identification accuracy from single-trial fMRI activity (Fig. 4e and Extended Data Fig. 10), demonstrating the potential applicability of our method to experiences that are difficult to reproduce, such as dreams^25^.

## Discussion

We have successfully generated descriptive text representing visual content experienced during perception and mental imagery by aligning semantic features of text with those linearly decoded from human brain activity. Our success is attributed to two key factors: the advancement of deep LMs that provide contextualized semantic representations similar to those in the human brain^28–33^ and our innovative text optimization method utilizing the MLM model for word-level optimization while efficiently constraining the search space. Together, these factors facilitate the direct translation of brain representations into text, resulting in faithful descriptions of visual semantic information in the brain. These descriptions were well structured, accurately capturing individual components and their relationships without using the language network, thus highlighting the explicit representation of fine-grained structured semantic information outside this network. Our method enables the intelligible interpretation of internal thoughts, demonstrating the feasibility of non-verbal, thought-based brain-to-text communication.

One distinct feature of our method compared to previous visual semantic decoding approaches is that it generates descriptions without enforcing linguistic coherence. Most prior methods, including those utilizing caption databases^17^ and non-linear image captioning models^18–20^, are inherently designed to produce linguistically structured outputs, limiting their capacity to probe for structured information explicitly represented in the brain. In contrast, as shown by the inability of decoders trained on word-order shuffled captions to generate coherent sequences (Extended Data Fig. 3c), our text optimization method (cf., Fig. 1b) permits the generation of incoherent outputs, particularly when the linguistic structure is deliberately removed from the training data. Yet, our method still produced coherent descriptions when decoding from the brain, progressively aligning more closely with brain representations—even from a non-informative initial state (i.e., <UNK> token; Fig. 2). This suggests that the coherence observed in the generated descriptions genuinely reflects structured information encoded within the brain. By elucidating the origins of the coherence, our method ensures a faithful decoding of contextualized semantic representations from the brain.

While our method generates linguistic outputs through brain decoding, it differs from previous language decoding attempts as it does not rely on brain activity associated with language production^2,4^ and perception^1,3^. Instead, we trained decoders on brain activity induced by non-linguistic visual stimuli to predict semantic features linked to the visual content of viewed videos. This approach enabled the generation of descriptions for both viewed and recalled content without involving the language network (Figs. 3 and 4).

Notably, all subjects in our study were non-native English speakers; nevertheless, our method proficiently generated text outputs in English. These results illustrate that our method can directly translate non-linguistic brain semantics into linguistic descriptions, regardless of the subject’s native language or language proficiency. Consequently, it can be applied to decode brain activity in non-linguistic subjects, including infants and animals, providing insights into how they develop the neural basis for processing complex visual semantics.

Moreover, by enabling the translation of non-verbal visual semantics in the brain into text, our method opens new communication channels in BMI applications, extending possibilities beyond traditional approaches for individuals with language or motor impairments. For instance, this approach could serve as an effective means of communication for individuals with aphasia, who struggle with language expression due to damage in language areas. Furthermore, our method complements visual-based BMI systems^55^ by providing an alternative communication pathway for individuals with conditions like amyotrophic lateral sclerosis (ALS), where degeneration of motor-related activity limits the effectiveness of motor-based BMIs. Thus, our method holds the potential to enhance communication and interaction in clinical and assistive settings.

The present study sheds light on the neural bases of structured visual semantics by examining the effects of ablating the language network and the posterior parts of category-selective regions during text generation analysis (Fig. 3). Excluding these regions did not substantially impact the quality of the generated descriptions, despite the broad distributions of semantic-feature-predictive voxels across the cortex, including within these regions—consistent with previous studies^44–47^. This outcome underscores the significance of brain areas outside these regions in representing structured visual semantics and aligns with research on visual representations involved in recognizing interactions and actions^11–16^. Our findings build on these studies by demonstrating that such representations are sufficient for constructing fine-grained, cohesive descriptions of both viewed and recalled content.

Furthermore, our encoding analysis, which contrasts models based on LM-derived contextual semantic features with visual DNN features related to object and action categories, revealed a functional boundary within the category-selective areas that separates posterior from anterior regions (Fig. 3b). Although this analysis specifically focused on neural representations linked to visual stimulus perception, this boundary coincides with known distinctions between visual and linguistic semantics (videos vs. audio stories)^56^ and between perceptual and mnemonic systems^57^. Consequently, our findings suggest an alternative perspective on this boundary: posterior regions may primarily support isolated semantic (or categorical) representations, while anterior regions integrate these into contextualized representations. The proximity of these anterior regions to language areas suggests they may play a bridging role, transforming non-verbal information into verbal expressions and connecting non-verbal and verbal semantics. Further research is needed to clarify how these anterior regions interact with the language network to achieve this integration.

While our method has shown the ability to generate descriptions that resemble captions rated by the subjects as highly consistent with their perceptions (Extended Data Fig. 5e), there remains potential for improvement in capturing the full spectrum of subjective experiences, particularly by refining the alignment and depth of captions annotated to video stimuli. Specifically, we relied on captions provided by independent annotators, which may not fully align with each subject’s unique perceptions, potentially affecting decoding performance. Although our use of rich annotations for each video (20 captions per video) likely mitigated some variability, training decoders on subjects’ own reports might yield even closer alignment. Furthermore, because we instructed annotators to focus on visual content rather than subjective aspects like emotional reactions^58^, the generated descriptions were predominantly concrete and rarely reflected abstract dimensions such as impressions and emotions^59^. With annotations that more accurately reflect and encompass various dimensions of subjective experience, our method may capture the content of a subject’s mind more comprehensively.

A limitation of our study is the use of natural videos originally sourced from the web^35^, which may restrict our ability to assess the generalizability of our method to atypical scenes (e.g., “a man bites a dog”). Notably, the word orders of the generated descriptions were successfully optimized to align with brain-decoded features (Fig. 2f), effectively capturing the relational information of the visual content (Fig. 2g). These results suggest that the generated descriptions faithfully reflect the structured visual semantics represented in the brain. However, it remains uncertain whether our method can generalize beyond common relational patterns or if it exhibits a bias toward typical structures. Future studies could address this limitation by incorporating controlled stimuli designed to depict contrasting or unusual relationships among elements^41,60^, allowing for a more rigorous assessment of our method’s generalization performance across a wider range of relational contexts.

A potential concern with our demonstration of generating mental content descriptions is that the verbal prompts used to cue the target videos may have influenced brain activity during the imagery period due to the slow hemodynamic response. During the preparation period, subjects might have started to recall the videos while reading these prompts, making it difficult to fully differentiate brain activity associated with text reading from that related to mental imagery. However, because our decoders were specifically trained on brain activity induced by non-linguistic visual stimuli, they prioritize semantic information directly linked to visual content over linguistic cues.

Furthermore, our analysis showed that descriptions generated during the imagery period were of higher quality than those from the preparation period (Extended Data Fig. 9c, d), suggesting that volitional mental imagery effectively recruited the neural representations necessary for accurate descriptions. Nonetheless, future investigations applying our method to spontaneous mental imagery (e.g., mind-wandering or dreaming) using subjects’ verbal reports as a reference would be necessary to clarify its ability to generate descriptions of mental content free from the influence of external stimuli.

The generation of mental content descriptions was successful even from single-trial fMRI activity of mental imagery (Fig. 4e and Extended Data Fig. 10). However, this success raises ethical concerns regarding potential invasions of privacy. Key issues include the risk of unintentionally disclosing primitive thoughts before individuals have chosen to verbalize them. Additionally, unwanted biases inherent in the LMs^61^ could distort the results within feasible optimization limits. Moreover, although requiring intensive data collection from willing participants may currently ensure consent^3^, advances in inter-individual alignment technology could reduce this requirement^62,63^.

Therefore, it is imperative to establish regulations that promote the ethical use of these technologies^64^ while ensuring explicit informed consent and safeguarding subjects’ mental privacy and autonomy in deciding which thoughts to disclose.

## Methods

### Subjects

Six healthy subjects (S1, male, aged 37–38; S2, female, aged 37–38; S3, male, aged 33– 34; S4, female, aged 35–36; S5, male, aged 29–30; and S6, male, aged 22–23) with normal or corrected-to-normal vision participated in the experiments. All were native Japanese speakers and non-native English speakers; S1–S5 were proficient in English, while S6 had limited proficiency. All subjects provided written informed consent, and the study protocol was approved by the Ethics Committee of NTT (R03-004 and R03-009). The sample size was determined based on prior fMRI studies with similar protocols^7,59^. Data from each subject were collected over multiple scanning sessions spanning approximately six months. Experimental parameters and analytical pipelines were determined from a preliminary experiment with S1, who was exposed to the same stimuli multiple times, potentially influencing their brain responses.

### Visual stimuli

Visual stimuli consisted of 2,196 short videos (all, 0.152 s to 90.1 s, mean = 6.61 s, median = 4.51 s; training, 0.152 s to 90.1 s, mean = 6.70 s, median = 4.62 s; test, 0.30 s to 20.1 s, mean = 3.94 s, median = 2.90 s) from a previous study^35^ (https://goo.gl/forms/XErJw9sBeyuOyp5Q2). These videos covered diverse content (objects, scenes, actions, and events) and were resized to fit a 16-degree visual angle, maintaining the original aspect ratio. They were presented at the center of a gray background without sound. Sixteen duplicates were excluded to avoid redundancy, resulting in the final set of 2,180 unique videos used in our experiment.

### Experimental design

We conducted two main experiments: a video presentation experiment and an imagery experiment (Extended Data Fig. 1). Visual stimuli were shown on an LCD monitor at the rear of the fMRI scanner, and audio stimuli were delivered via S14 earphones (Sensimetrics). Each experimental session lasted up to 2 hours. Subjects were given adequate time for rest between runs (every 8–10 minutes) and could take a break or stop the experiment at any time. The total duration for both experiments was approximately 17.1 hours.

### Video presentation experiment

The video presentation experiment included training sessions (60 runs) and test sessions (10 runs). Each run had 36 or 37 stimulus blocks and 3 or 4 randomly inserted evaluation blocks, averaging 695.2 s per run. Subjects spent approximately 11.8 hours on training sessions and 1.8 hours on test sessions. In each stimulus block, videos shorter than 10 s were repeated until the total duration exceeded 10 s. Videos longer than 10 s were presented once, followed by less than 1 s of rest, making the block duration divisible by 1 s (TR). Subjects viewed the stimuli without fixation to recognize details.

In each evaluation block, five descriptions (both in English and translated into Japanese) depicting the visual contents of videos were presented, and subjects were then asked to rate how consistent each description was with what they had perceived in the preceding stimulus block using two button boxes. The descriptions were taken from twenty captions for the preceding video, with occasional ones from other videos. Subjects rated the descriptions on a five-point scale or marked them with an “x” if they were unrelated (with higher scores indicating closer alignment with their perception; random initial score). Subjects completed the ratings at their own pace and proceeded to the next block by selecting “Proceed?” and pressing a button.

Each block was followed by a 2-s rest period, with 32-s and 8-s rest periods at the beginning and end of each run, respectively. During the rest periods, the subjects were instructed to maintain fixation on a central spot, which consisted of a bull’s eye and crosshairs, in order to keep their attention focused on the screen.

In the training session, 2,180 unique videos were each presented once in a pseudo-randomized order. This order remained consistent for all subjects. Evaluated descriptions were also consistent for all subjects. In the test session, 72 videos from the last two runs of the training session were each presented five times, divided between two runs, and shown in a pseudo-random order within each run (see Methods “MRI data preprocessing” for the data handling of the overlapping data in the training session).

### Imagery experiment

The imagery experiment comprised 30 runs, each with 12 trials consisting of a preparation block, an imagery block, a video presentation block, and an evaluation block. Subjects were required to engage recall-based visually imagery of one of the 72 videos presented during the test session of the video presentation experiment. Each run averaged 434.6 s, totaling approximately 3.6 hours per subject.

During a preparation block, a verbal description (both in English and translated into Japanese) of a target video was presented to subjects to prompt them to prepare mental imagery of the visual content of the target. The descriptions were selected from the set of twenty captions collected for each video and were consistent across trials and subjects. Subjects were encouraged to imagine all details, even those not explicitly described, to mimic the video presentation experiment. The description served as a guide to vividly imagine the complete visual content of the target. Subjects pressed a button when ready, and a beep sound with less than 1 s of rest signaled the start of the imagery period, aligning with the TR.

During the imagery period, subjects recalled the visual content of the target video with their eyes closed as if actually watching it. The imagery block duration matched the stimulus block duration in the video presentation experiment (repeatedly recalling for videos shorter than 10 s, once for videos longer than 10 s), with an additional 2 s to ensure full replay. A beep signaled the end of the imagery period, prompting subjects to open their eyes.

After the imagery block, the target video was presented as in the video presentation experiment, allowing subjects to compare their mental imagery with the actual video. This was followed by two 3-s blocks where subjects rated the accuracy and vividness of their imagery on a five-point scale. Subjects adjusted the score from its random initial setting using a button box in their right hand.

Each imagery, stimulus, and evaluation block was followed by a 2-s rest period, with 32- and 8-s rest periods at the beginning and end of each run. Subjects maintained fixation on a central spot, as in the video presentation experiment.

Before the imagery experiment, subjects practiced associating each target video with its verbal description, viewing the pairs during inter-run rest periods to aid memory. The 72 videos were randomly distributed among six runs, with each set of six runs containing all videos in a pseudo-random order.

### Retinotopy and functional localizer experiments

In addition to the main experiments, we conducted a retinotopy experiment and three functional localizer experiments (visual category, MT+, and language area localizers) to delineate visual areas and localize regions of interest (ROIs).

#### Retinotopy

We followed the Human Connectome Project 7T Retinotopy Dataset protocol^65^, using dynamic colorful textures through moving apertures (wedge, ring, bar) in eight 300-s runs. This identified retinotopic maps (V1, V2, V3, V3A, V3B, hV4, and V7) on cortical surfaces using *fsfast* retinotopy analysis (*Freesurfer*^66^) and population receptive field analysis (code is available at https://kendrickkay.net/analyzePRF/)^67^.

#### Visual category localizer

Following the fLoc protocol^68^ (stimuli and code are available at http://vpnl.stanford.edu/fLoc/), we presented images from word, body, face, place, and object categories in eight 300-s runs (48 blocks each, 6 s per stimulus or blank, with 6-s initial and final rests). Additional intact and scrambled object conditions were included to localize object-selective areas (original images are available at tarrlab, https://sites.google.com/andrew.cmu.edu/tarrlab/stimuli). The contrasts between word/body/face/place and others (from these four categories) were used to define visual category-selective areas (word: visual word form area [VWFA], occipital word form area [OWFA]; body: extrastriate body area [EBA], fusiform body area [FBA]; face: fusiform face area [FFA]; occipital face area [OFA]; place: parahippocampal place area [PPA], occipital place area [OPA], medial place area [MPA] consisted of the retrosplenial cortex [RSC] and parieto-occipital sulcus [POS]). The contrast between intact and scrambled objects was used to define an object-selective area (lateral occipital complex [LOC]).

#### MT+ localizer

Following Tootell et al. (1995)^69^, we presented random dot stimuli in three conditions (moving, dynamic, static) in four 232-s runs (13 blocks each, 12 s per stimulus, with 12-s initial and final rests). The contrast between moving and dynamic/static conditions defined the visual motion area MT+.

#### Language area localizer

We followed protocols by Fedorenko et al. (2010)^70^ and Scott et al. (2017)^71^, modifying to include both visual and auditory stimuli and to use Japanese stimuli in each of eight 358-s runs (19 blocks each, 18 s per stimulus or 14-s blank, with 14-s initial and final rests). Subjects read sentences or nonword sequences and listened to intact or degraded auditory passages. The contrasts between sentence/intact and nonword/degraded conditions defined language-sensitive areas in the temporal and frontal cortices.

Voxels from V1, V2, and V3 were combined as the lower visual cortex (LVC); voxels from VWFA, OWFA, EBA, FBA, FFA, OFA, PPA, OPA, MPA, LOC, and MT+ were combined as the higher visual cortex (HVC); voxels from temporal and frontal language areas were combined as the language network. Overlapping voxels with LVC were excluded from HVC.

### MRI acquisition

MRI data were collected using a 3.0-Tesla Siemens MAGNETOM Prisma scanner located at the WPI-IRCN Human fMRI Core, the University of Tokyo Institutes for Advanced Studies. An interleaved T2*-weighted gradient-echo echo planar imaging (EPI) scan was performed to acquire functional images covering the entire brain (TR, 1000 ms; TE, 30 ms; flip angle, 65 deg; FOV, 192 × 192 mm; voxel size, 2 × 2 × 2 mm; slice gap, 0 mm; number of slices, 72; multiband factor, 6). T1-weighted (T1w) magnetization-prepared rapid acquisition gradient-echo (MP-RAGE) fine-structural images of the entire head were also acquired (TR, 2000 ms; TE, 1.97 ms; TI, 900 ms; flip angle, 10 deg; FOV, 256 × 256 mm; voxel size, 1.0 × 1.0 × 1.0 mm).

### MRI data preprocessing

For anatomical data, we first used *SPM12* to preprocess each of the T1w anatomical images of individual subjects for bias-field correction and for re-defining its origin and orientation to be set on the anterior commissure and the anterior commissure - posterior commissure (AC-PC) line, respectively. Cortical surface meshes were generated from the processed T1w images using *Freesurfer* (version 7.3.2)^66^ with manual corrections for anatomical segmentations. Analytical results were visualized on flattened cortical surfaces, created by making relaxation cuts in each hemisphere, with functional data aligned and projected using *Pycortex*^72^.

For the functional data from each run, we performed the MRI data preprocessing through the pipeline provided by *fMRIPrep* (version 20.2.7)^73^. First, a BOLD reference image was generated using a custom methodology of *fMRIPrep*. A field map (B0-nonuniformity map) estimated based on a phase-difference map was used to estimate susceptibility distortion and to correct the BOLD reference for a more accurate co-registration with the anatomical reference. The BOLD reference was then co-registered to the T1w reference using *bbregister* (FreeSurfer; version 7.3.2), which implements boundary-based registration^74^. BOLD runs were slice-time corrected using 3dTshift from AFNI 20160207^75^, and the BOLD time series were resampled onto their original, native space (2 × 2 × 2 mm voxels) by applying a single, composite transform to correct for head-motion and susceptibility distortions using antsApplyTransforms from ANTs (version 2.3.3) with Lanczos interpolation.

To create data samples, we first discarded the first 8-s scans of the preprocessed BOLD signals from each run to avoid MRI scanner instability. We then regressed out nuisance parameters from each voxel amplitude for each run, including a constant baseline, a linear trend, and 24 head-motion parameters (three rotations, three translations, their temporal derivatives, and quadratic terms) and 12 global signals (mean amplitudes within cerebrospinal fluid, white matter, gray matter, their temporal derivatives, and quadratic terms). The data samples were temporally shifted by 4 s to account for hemodynamic delays, despiked to reduce extreme values (beyond ± 3 SD for each run), and averaged within each stimulus and imagery block. Finally, each voxel’s amplitude was z-scored within each run to eliminate potential non-stationarities and scanner-specific biases.

For data from the training session of the video presentation experiment (training perception data), we discarded samples from the last two runs, in which videos used in the test session and the imagery experiment were presented, to ensure generalization to new stimuli. For test data from the video presentation experiment (test perception data) and the imagery experiment (test imagery data), we averaged samples of identical video clips (5 repetitions) to increase the signal-to-noise ratio of the fMRI signals unless otherwise stated.

### Caption annotation for visual stimuli

We used Amazon Mechanical Turk (AMT) to collect written captions for stimulus video clips, following the procedure used for Microsoft COCO Captions^76^. Multiple workers viewed each video to provide a detailed sentence (over eight words) describing the visual content. The captions were manually checked for quality and proofread with the assistance of ChatGPT (GPT-3.5; https://chat.openai.com/; prompt: “Proofread the following:”) to correct typos and remove incorrect or unintelligible sentences. We collected twenty unique captions per video, matching the number in the MSR-VTT dataset^77^. The collected video captions are available from our repository (https://github.com/horikawa-t/MindCaptioning).

### Feature computation by deep neural network models

We used deep neural network models pre-trained for language or vision tasks to compute semantic and visual features. To mitigate biases arising from baseline differences across model units, we applied *z*-score normalization to the feature values using means and standard deviations estimated from respective training data for each analysis.

#### Semantic features

To extract semantic features from video captions, we used 42 pre-trained language models (LMs; available at Hugging Face’s Transformers library, version 4.30.2)^78^. These models cover a range of architectures (e.g., BERT and GPT2) and sizes (e.g., base, large, and xlarge). Each input sequence was tokenized and processed by an LM to produce vector embeddings for each token across multiple layers. Following Reimers & Gurevych (2019)^79^, we averaged the embeddings over tokens, excluding special tokens (e.g., <CLS> token), in each layer. The averaged embeddings from multiple layers were used as semantic features for the input sequence. For each video, we computed semantic features for 20 annotated captions and averaged them to construct the final semantic features for the video.

The 42 LMs used in the present study were based on the following model families: BERT^34^ (*33*), RoBERTa^37^, DeBERTa^36^, ALBERT^80^, OpenAI-GPT^81^, GPT2^82^, Sentence GPT^83^, XLnet^84^, DistilBERT^85^, DistilGPT2^78^, T5^86^, BART^87^, CTRL^88^, XLM^89^, XLM-RoBERTa^90^, ELECTRA^91^, CLIP^92^ text encoder. See Extended Data Fig. 6 for the full list of the LMs.

We hypothesized that an LM closely aligned with the human brain will provide more effective intermediate representations for translating visual semantic information in the brain into text. We thus performed a cross-validation encoding analysis using semantic features from each of the 42 LMs within the training perception data. Based on the results of the validation analysis (Extended Data Fig. 2a), we selected the DeBERTa-large model, which demonstrated the highest performance.

To construct semantic features without structured semantic information (Extended Data Fig. 3c), we used captions with randomly shuffled word orders. For each caption, we created up to 1,000 word-order shuffled variants, computed their semantic features, and averaged these features across all the variants. These features were then averaged across twenty captions for each video to obtain semantic features for each video.

#### Vision model features

To extract visual features from video stimuli, we used a TimeSformer model^53^ (model and code are available at https://github.com/facebookresearch/TimeSformer) pre-trained for object and action recognition using *ImageNet*^93^ and Kinetics-400^94^. This model has demonstrated high performance in predicting video-induced brain activity^95^. For each video, we resized its spatial size to 224 pixels while preserving the aspect ratio, and selected frames at intervals of 32. If the video had fewer than eight temporal positions at 32-frame intervals, we uniformly selected eight frames to cover the entire video length. We computed feature vectors for each layer from these frames and averaged them over the spatial dimension within each layer. This procedure was repeated for three spatial crops (left-center-right or top-center-bottom). Finally, we averaged the visual features over the temporal dimension and the three spatial positions, resulting in 768 features for each of the 12 layers.

### Voxel-wise encoding modeling analysis

We performed voxel-wise encoding modeling analysis by constructing encoding models that predict signal amplitudes of individual voxels from a feature vector in each model layer using the L2 regularized linear regression algorithm (ridge regression). The analysis was performed using both cross-validation and generalization approaches. In the cross-validation analysis, we used 6-fold cross-validation on the training perception data (58 runs divided into five sets of 10 runs and one set of 8 runs). In the generalization analysis, we trained models on all of the training perception data and tested them on the test perception data (Extended Data Fig. 6). We evaluated the model performance of each voxel by calculating Pearson correlation coefficients between measured and predicted brain activities of that voxel.

The regularization parameters of ridge regression were determined separately for each layer, model, and subject by considering the performance of all voxels on the respective training data. Models for individual voxels were trained using ten possible regularization coefficients (log spaced between 10 and 10000). The regularization parameters that produced the maximal model performance (mean correlation coefficients averaged across all voxels) on the training data were used for predictions on the test data. In the cross-validation analysis, we used a 5-fold cross-validation (inner loop) nested within a 6-fold cross-validation (outer loop). In the generalization analysis, we used 6-fold cross-validation on the training perception data.

For each model, predictions from multiple layers were integrated by selecting the best layer for each voxel based on model performance in the training data. In each fold of the cross-validation analysis, we determined the best layer per voxel from the 5-fold nested cross-validation loops (cf., Fig. 3f). We aggregated predictions from these best layers for each left-out set to construct final predictions for all data samples. In the generalization analysis, the best layer per voxel was determined from the 6-fold cross-validation on the entire training perception data, and predictions from the best layers were used for the test perception data.

Encoding accuracies of semantic and visual models were compared using slope angles of the best linear fit estimated by Deming regression^96^, which accounts for observation errors on both axes. The slopes were converted to angles, then subtracted from 45 degrees to obtain deviations from parity (Fig. 3e).

### Feature decoding analysis

We performed feature decoding analysis by constructing a set of L2 regularized linear regression models (decoders) that predict feature values for each layer of each model from fMRI activity patterns (one decoder for each model unit). Both cross-validation and generalization analyses were performed to produce predictions for all training perception data samples as well as the test perception and imagery data samples. Decoders were trained using whole-brain fMRI voxel patterns (unless otherwise stated), selecting up to 50,000 voxels best predicted by the target feature set in cross-validation (or nested cross-validation) encoding analysis on the respective training data. Performance was evaluated by calculating Pearson correlation coefficients between feature values computed by an LM and predicted from the brain for each model unit.

In the cross-validation analysis, we used 6-fold cross-validation on the training perception data, with model training and the regularization parameter estimation conducted using a 5-fold cross-validation procedure nested within the 6-fold cross-validation, similar to the encoding analysis. In the generalization analysis, models were trained on all the training perception data and tested on the test perception and test imagery data, with regularization parameters determined using a 6-fold cross-validation within the entire training perception data. Regularization parameters of the ridge regression models were optimized based on model performance on the respective training data, with coefficients estimated separately for each layer, model, and subject considering the performance of all units.

### Text generation analysis

To generate descriptive text based on a set of semantic features (target features), we conducted an iterative optimization of descriptions, in which semantic features of candidate descriptions were progressively aligned with target features through the iterative replacement and interpolation of tokens (referred to as “words” for simplicity) within the candidate descriptions. Each step of the optimization process consisted of three stages: masking, unmasking, and candidate selection.

In the masking stage, for each of the current candidate descriptions (e.g., “metal shapes”), we first generated an exhaustive list of masked candidates by replacing each word or a sequence of words (three words at the maximum) with a mask token (e.g., “<MASK> shapes,” “metal <MASK>,” and “<MASK>”) or interpolating a mask token between words or at the top or bottom of the description (e.g., “metal <MASK> shapes,” “<MASK> metal shapes,” and “metal shapes <MASK>”). This masking procedure was repeatedly applied on the newly generated masked candidates (two times at the maximum) to generate masked candidates with multiple masks (e.g., “<MASK> <MASK> shapes” and “metal <MASK> <MASK>”). From all the generated masked candidates for each original candidate, we randomly selected five masked candidates to be processed in the next unmasking stage.

In the unmasking stage, we used an LM pre-trained for masked language modeling (MLM model; not necessarily the same as the model used for the feature computation) to generate alternative words to fill in the masks in the masked candidates within the context of surrounding words. For each mask within a masked candidate description, we generated five alternative words for the mask to create five new candidates by random sampling from a categorical distribution likelihood estimated by the MLM model. In cases where a masked candidate had multiple masks, we processed the masks sequentially from the top until all masks were updated. These procedures yielded five new candidates from one masked candidate. In the main analysis, we used the pre-trained RoBERTa-large model (vocabulary size, 50265; subword segmentation based on byte pair encoding) for the guide of the text generation because this model consistently demonstrated stable performance in optimizing descriptions in a validation analysis (Extended Data Fig. 2b).

In the candidate selection stage, we computed semantic features of all new and original candidate descriptions using an LM from which the target features originated. We then calculated Pearson correlation coefficients between those candidate features and target features for all layers and averaged those correlations over layers to score the candidate descriptions. To enhance the conciseness of generated descriptions, reduce the computational cost of handling long descriptions, and avoid overfitting to noises on brain-decoded features, we added an exponential penalty to the length of candidate descriptions as described by 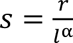, where *s* is the similarity score used to rank candidates, *r* is the mean correlation coefficient between candidate features and target features averaged across layers, *l* is the length (or the number of tokens) of the candidate, and *α* is the parameter for the length penalty. We chose the parameter *α* = 0.1 based on the validation analyses using six possible penalty parameters (0, 0.05, 0.1, 0.15, 0.2, and 0.25) with a randomly selected subset of 50 samples from the training perception data (Extended Data Fig. 2c–e). After computing similarity scores for all candidates, we ranked them and selected the top five candidates to proceed with further optimization.

During each step of the optimization process, the maximum search width was 130 (five new candidates for each of the five masked candidates derived from five original candidates, in addition to the five original candidates themselves).

We repeated these optimization stages 100 times, and the obtained description was taken to be the text describing the semantic information represented in the target features, or the brain. Since the masking and unmasking stages involve randomness in the optimization process, to avoid local optima we repeated the same process five times for each data sample to select the description showing the highest similarity scores with the target features as the final prediction. For all the analysis, we began the optimization process from a non-informative initial state (i.e., unknown token, <UNK> for the tokenizer of the RoBERTa-large) to avoid incorporating any prior assumptions for description generation.

Unlike autoregressive LMs with causal attention often used for linguistic information decoding^3,4^, MLM models with bidirectional attention have the significant advantage of incorporating contextual information from all surrounding words^34^. This characteristic makes our optimization process more suitable for decoding visual information, which lacks specific directionality, in contrast to linguistic information.

### Database-search-based description prediction analysis

The main strengths of our method are its flexibility to optimize descriptions at the word (or token) level and its ability to generate word sequences that do not currently exist in the databases. To assess its effectiveness, we used the database-search (DB-search) method^17^ to set a baseline performance for description prediction from the brain. This analysis involved searching for captions with the highest feature correlations with target brain-decoded features from large databases of image and video captions [MS-COCO^97^; GCC^98^; MSR-VTT^99^]. We computed semantic features for all captions (∼4.1M) using the DeBERTa-large model. For each fMRI data sample, we predicted semantic features at multiple layers using trained decoders and computed correlation coefficients between the decoded features and features computed from all database captions. The caption with the highest mean correlations across all layers was selected as the prediction for the sample. Note that the brain-decoded features used in this analysis were the same as those used in the main analysis (i.e., mind captioning; e.g., Fig. 2).

### Evaluation of the similarity between generated descriptions and references

We used multiple metrics to evaluate the similarity of generated descriptions to reference captions annotated to the viewed or recalled videos. These metrics include Pearson correlation (feature correlation), BLEU, METEOR, ROUGE-L, CIDEr, and three variants of BERTScore. For each predicted description, we computed scores for each metric against twenty reference captions of the corresponding video, selecting the highest score as the final score. BLEU and METEOR were computed using the NLTK toolbox^100^, while ROUGE-L and CIDEr were computed using code from https://github.com/salaniz/pycocoevalcap. BERTScore was computed using code from https://github.com/Tiiiger/bert_score.

#### Feature correlation

We defined the feature correlation as the mean of Pearson correlation coefficients between semantic features of a reference caption or target features and those of a generated description averaged across multiple layers. We computed semantic features using the same LM as in the decoding analysis, unless otherwise stated. Feature correlation was used in all analyses where the similarity metric was not explicitly specified.

#### BLEU

BLEU computes the precision scores by comparing predicted n-grams with reference captions while considering a brevity penalty. We used the 4-gram variant (BLEU-4) with a smoothing method^101^.

#### METEOR

METEOR computes scores based on unigram matching in predicted and reference sequences using precision and recall while considering word variations like stemming and synonymy.

#### ROUGE-L

ROUGE-L computes scores by emphasizing recall, measuring the overlap in words between the prediction and reference, and primarily focusing on their longest shared sequence.

#### CIDEr

CIDEr measures the similarity between predicted descriptions and references through n-gram-based comparison while considering consensus across multiple references.

#### BERTScore

BERTScore computes similarity using contextualized embeddings from individual tokens by a bidirectional transformer LM. We used the 17th layer of the RoBERTa-large model with baseline rescaling to compute scores (default of the official implementation) and evaluated three variants: P (precision), R (recall), and F1. To evaluate the token-wise precision of generated descriptions, we used the precision (P) weighted by the inverse of document frequency (IDF) estimated from captions in multiple databases (MSCOCO, GCC, and MSR-VTT) and captions collected in this study. The IDF-weighted P was only used to highlight tokens with high precision in generated descriptions, but not used in quantitative evaluations.

Based on these metrics, we evaluated the discriminability of the generated descriptions. For each description, we calculated similarity scores between the generated description and reference captions of the corresponding viewed or recalled videos (correct) and irrelevant videos (incorrect; *n* = 2,179). Discriminability was defined as the difference between the score for the correct video and the mean score averaged across all incorrect videos.

### Video identification analysis

To evaluate description generation performance, we performed a video identification analysis based on the similarity between generated descriptions and reference captions using multiple metrics. For each data sample, we computed similarity scores between the generated description and all reference captions of all videos using a specific metric. We compared the similarity scores to the correct reference captions (20 captions for the target video) with those to incorrect reference captions (43,580 captions for 2,179 irrelevant videos). For the analysis with feature correlation and BERTScore, we used mean similarity scores averaged across multiple captions per video. For BLEU, METEOR, ROUGE-L, and CIDEr, we used the highest matching scores among multiple captions per video. The analysis was conducted with varying numbers of candidates, ranging from 2 (chance level, 50%) to 100 (chance level, 1%), selecting the video with the highest similarity score as the prediction. Accuracy was defined by the proportions of correct video identification. For two candidates (one correct and the other incorrect), we performed identification for all combinations of correct and incorrect candidates. For more than two candidates, we randomly selected incorrect candidates, repeating the analysis 100 times to estimate the mean accuracy averaged across repetitions.

### Evaluation of the fidelity of the relational information in generated descriptions

To assess whether the generated descriptions faithfully represented visual relationships among individual components in viewed or recalled videos (e.g., differences between “a bird eats a snake” and “a snake eats a bird”)^41^, we evaluated the effect of shuffling word orders of generated descriptions on discriminability and video identification performance. We randomly shuffled the word order of each generated description to create word-shuffled variants. The shuffling was performed at the word level, not the token level, to maintain minimal coherence within individual words. For each original description, we created up to 1,000 shuffled variants by shuffling all words or only nouns, excluding descriptions with only one noun from the noun-shuffling analysis. Nouns were identified using part-of-speech tagging with *spaCy*^102^ (version 2.2.4). We then computed semantic features for these shuffled descriptions.

If generated descriptions accurately capture visual relations among individual components in videos, we should observe higher similarity to correct captions with the original descriptions compared to the shuffled ones while maintaining differences with irrelevant captions. To examine this, we computed feature correlation scores between reference captions and both original and shuffled descriptions to examine if the original descriptions exhibited higher discriminability (e.g., Fig. 2f right). Given the varying levels of sentence structure disruption among the shuffled descriptions, we also conducted the same analysis using the least disrupted shuffled sentences to rigorously assess the superiority of the original descriptions. Specifically, we selected the shuffled sentence with the highest pseudo-log-likelihood score—a fluency (or linguistic acceptability) metric computed by MLM scoring^42^—from 1,000 shuffled variants for each generated description (e.g., Extended Data Fig. 4f). We employed an adapted metric proposed by Kauf & Ivanova (2023)^43^. Additionally, to quantify the impact of shuffling on identification accuracy, we performed video identification analysis using both original and shuffled descriptions (e.g., Fig. 2f left).

We also examined if the word order of generated descriptions was unduly influenced by the MLM model used to support the text generation, potentially diverging from the information represented in the brain. We reasoned that if a generated description faithfully matches the brain representation and contains semantic information uniquely depicted by the generated word order—rather than alternative arrangements of the same words—then brain-decoded features should exhibit higher similarity to features of the generated description than features of shuffled variants. To test this, we computed feature correlation scores between target brain-decoded features and both original and shuffled descriptions to see if the original exhibited greater scores. To consider the degree of meaning changes introduced by shuffling, we also computed feature correlation scores between original and shuffled descriptions, defining correlation distance as one minus the feature correlation (Fig. 2g and Extended Data Fig. 4g).

### Evaluation of the diversity in generated descriptions

To evaluate the diversity of generated descriptions (Extended Data Fig. 5c, d), we used *Self-BLEU*^103^—a metric assessing the diversity of the generated text data—to compute the sentence (dis)similarity either across subjects for each video or across videos for each subject. For each video/subject, we can compute a BLEU score by regarding one description from a subject/video as a hypothesis and descriptions from the other subjects/videos as references. To evaluate the diversity across subjects/videos, we computed BLEU scores for every generated description from different subjects/videos and defined the average BLEU scores across subjects/videos as the Self-BLEU of the video/subject, respectively. A higher Self-BLEU score indicates less diversity in the descriptions generated for the video/subject. In this analysis, we used the same analytical settings as we did when using the BLEU for the similarity evaluation (BLEU-4 with smoothing).

### Evaluation of the consistency between generated descriptions and subjective perception

The reference captions in this study were collected from subjects in an independent online experiment, not from our fMRI subjects. Therefore, not all captions may precisely match the subjective perception of our fMRI subjects, even though they were used as “correct” references in the evaluation. To examine if the generated descriptions from video-induced brain activity were consistent with the subjective perceptions of individual subjects, we investigated the relationship between subjective ratings from the video presentation experiment and the similarity of generated descriptions to evaluated captions. We focused on 212 videos evaluated during the training perception data collection, analyzing descriptions generated from the cross-validation decoding analysis using feature correlations as the similarity metric. For each generated description, we computed feature correlations against five evaluated captions, yielding 1060 scores per subject. These scores were classified according to individual ratings to explore if higher-rated captions had higher scores. Additionally, we calculated Pearson correlation coefficients between the feature correlation scores and subjective ratings to determine the presence of positive correlations (Extended Data Fig. 5e).

### Statistics and reproducibility

Statistical analysis was performed individually unless otherwise stated, with results from six subjects considered replications^104^. We reported quantitative results for each subject and averages across subjects, except for the validation analysis (Extended Data Fig. 2c–e). To account for multiple comparisons, we used the Benjamini-Hochberg method^105^ to control the false discovery rate (FDR) and provided this information where applicable.

Statistical significance between results of pre-trained and untrained MLM models was tested using a one-tailed Wilcoxon signed-rank test on feature correlations of generated descriptions after 100 optimization iterations (*n* = 72; Extended Data Fig. 4a).

Discriminability based on generated descriptions was also tested using a one-tailed Wilcoxon signed-rank test (*n* = 72; e.g., Extended Data Fig. 4b). The effect size of discriminability (e.g., Fig. 2d) was computed by calculating similarity scores between a generated description and both sets of correct (*n* = 20) and incorrect captions (*n* = 43,580). These scores were averaged separately for each set. These averaged scores for correct and incorrect sets from all test samples (*n* = 72) were used to estimate means and standard deviations for computing Cohen’s *d* for discriminability.

Video identification analysis results were presented with 95% C.I. across samples (*n* = 72) to determine if the mean accuracy exceeded the chance level (e.g., Fig. 2e).

We used one-tailed Wilcoxon signed-rank tests to evaluate the impact of word-order shuffling on discriminability (*n* = 72; e.g., Fig. 2f) and the diversity differences in generated descriptions (*n* = 72 for self-BLEU across subjects; *n* = 6 for self-BLEU across videos; Extended Data Fig. 5c, d).

Correlations between discriminability and encoding accuracy across multiple LMs were evaluated using one-tailed *t*-tests after Fisher’s z transform (*n* = 42; Extended Data Fig. 6b). The correlation between layer depth and both discriminability and its drop caused by shuffling were tested using a one-tailed *t*-test (Extended Data Fig. 7).

To evaluate the similarity between the generated descriptions and rated captions, we pooled results from all six subjects to ensure sufficient data samples to detect differences, while also accounting for variations in the number of samples across rating levels and subjects (Extended Data Fig. 5e). Differences in feature correlations across subjective ratings were tested using one-tailed *t*-tests after Fisher’s z transform.

Correlation between ratings and feature correlations between generated descriptions and rated captions were tested using a one-tailed *t*-test after applying Fisher’s z transform (*n* = 6,360). Interactions between text generation methods and ratings were tested using ANOVA.

For encoding analysis, the correlation between measured and predicted fMRI signals for each voxel was tested using a one-tailed *t*-test after Fisher’s z transform (*n* = 2,108; Fig. 3a, b). Mean encoding accuracy within each brain area was presented with a 95% C.I. across voxels (Fig. 3d). Comparisons between semantic and visual encoding models were based on the slope angles of linear fits, converted to deviations from parity (Fig. 3e). Statistical significance of differences in the best layers exhibiting highest encoding accuracy was tested using a one-tailed Wilcoxon rank-sum test (Fig. 3f).

## Data availability

The experimental data that support the findings of this study are available from open data repositories (raw data in OpenNeuro: https://doi.org/10.18112/openneuro.ds005191.v1.0.2; preprocessed data in figshare: https://doi.org/10.6084/m9.figshare.25808179).

## Code availability

The code that supports the findings of this study is available from our repository (https://github.com/horikawa-t/MindCaptioning).

## Acknowledgements

The author thanks Yuki Honda for assistance with MRI data collection and cleaning caption data, Scinob Kuroki, Shimpei Yamagishi, Hiromi Narimatsu, and Yuta Suzuki for assistance with scanner operation, Mitsuaki Tsukamoto, Misato Tanaka, and Yukiyasu Kamitani for assistance with preliminary investigations, and Shogo Kajimura for discussions. We appreciate the support of the WPI-IRCN Human fMRI Core, the University of Tokyo Institutes for Advanced Studies. This research was supported by grants from JST PRESTO Grant Number JPMJPR185B Japan and JSPS KAKENHI Grant Number JP21H03536.

## Author contributions

Conceptualization, T.H.; Methodology, T.H.; Validation, T.H.; Formal Analysis, T.H.; Investigation, T.H.; Resources, T.H.; Writing – Original Draft, T.H.; Visualization, T.H.; Funding Acquisition, T.H..

## Competing interests

The author declares no competing interests.

## Additional information

**Supplementary Information** is available for this paper.

**Correspondence and requests for materials** should be addressed to Tomoyasu Horikawa.

## Extended Data

Extended Data Fig. 1 | Overview of experiments

Extended Data Fig. 2 | Validation of model and parameter determinations Extended Data Fig. 3 | Examples of generated descriptions

Extended Data Fig. 4 | Text generation performance of viewed content for individual subjects

Extended Data Fig. 5 | Comparison with database-search method

Extended Data Fig. 6 | Relations between encoding accuracy and text generation performance of multiple LMs

Extended Data Fig. 7 | Effects of shuffling on discriminability with varying numbers of layers

Extended Data Fig. 8 | Voxel-wise encoding model performance

Extended Data Fig. 9 | Text generation performance of recalled content for individual subjects

Extended Data Fig. 10 | Performance of text generation with different numbers of averaged samples

[Note: In Extended Data Figs 3 and 10, to comply with bioRxiv’s policy on displaying human faces, frames containing real human faces in this figure (framed by gray) have been replaced with images synthesized by DALL·E3 (https://openai.com/dall-e-3), based on captions annotated by humans for the video.]

**Extended Data Fig. 1.**
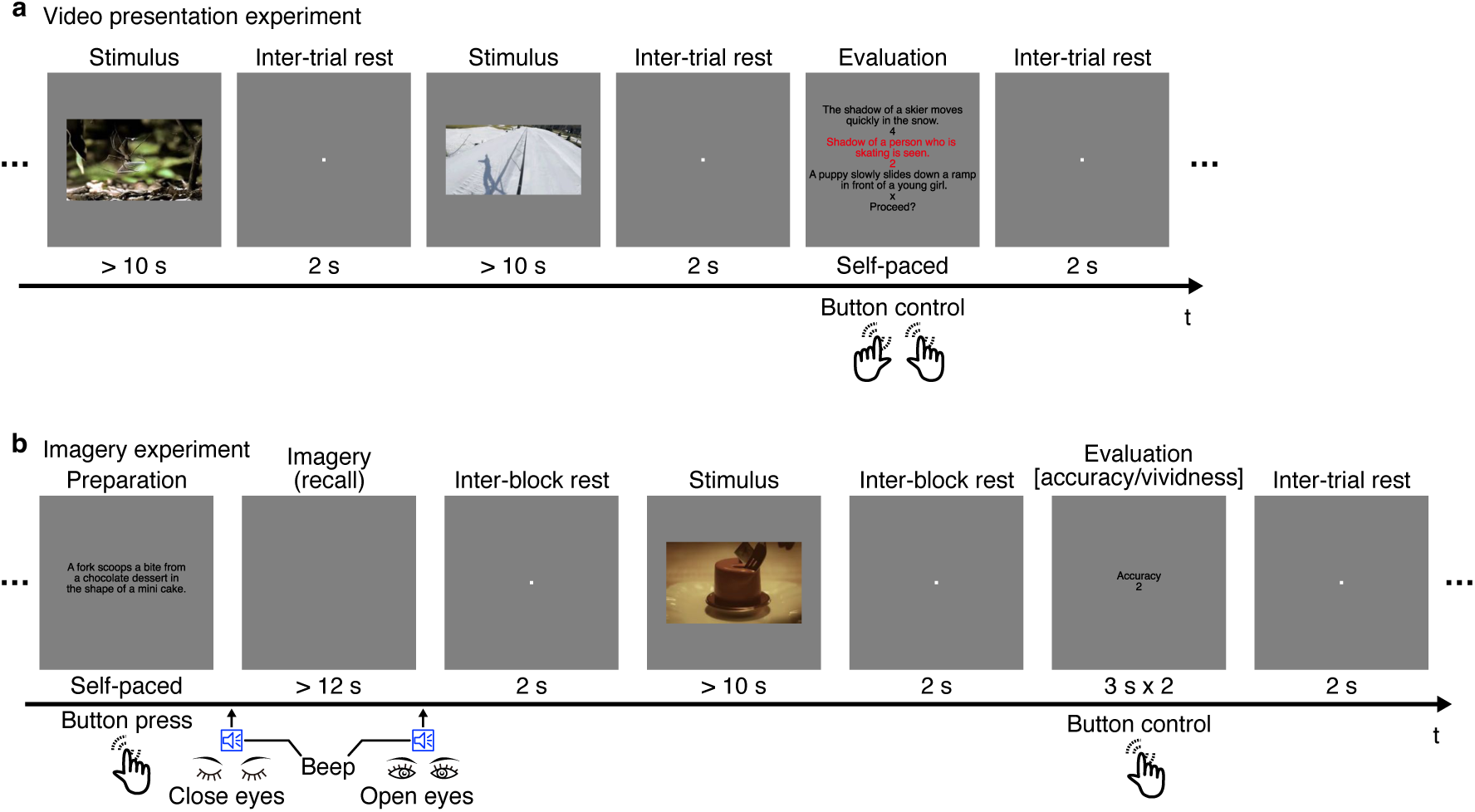
Overview of experiments. We conducted two types of experiments. **a**, Video presentation experiment (training and test sessions). Subjects were allowed to view videos without fixation. Occasionally, we presented five descriptions depicting the visual content of the preceding video to the subjects, asking them to rate the consistency between each description and their subjective perception. We used these ratings to evaluate the consistency between descriptions generated from the brain and the subjective perception of each individual (Extended Data Fig. 5e). **b**, Imagery experiment. Subjects were well-trained to associate descriptions and videos before the experiment. They were required to visually imagine (recall) a video based on a description of the target video with their eyes closed according to beeps. Each imagery block was followed by a stimulus block, during which the target video was presented to allow the subjects to confirm the validity of their imagery. An evaluation block followed, asking the subjects to evaluate the accuracy and vividness of their imagery. Data samples were constructed by averaging fMRI volumes during each stimulus/imagery block. Training data contained samples for 2,108 videos, each presented once. Test and imagery data contained samples for the same 72 videos (not used during training), averaged across five repetitions.

**Extended Data Fig. 2.**
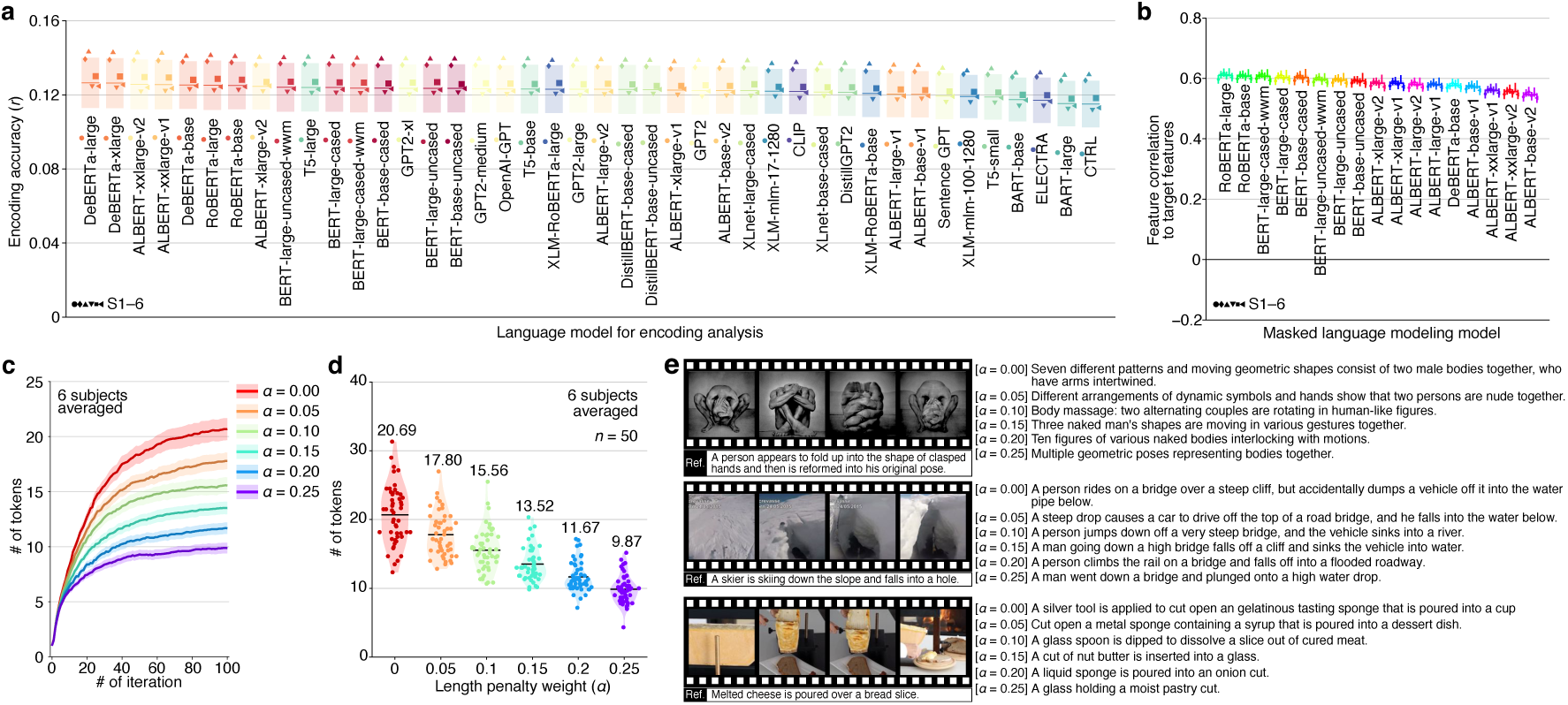
Validation of model and parameter determinations. Validation analyses were performed using cross-validation within the training perception data to determine LMs used in feature decoding and text generation analyses and a length penalty parameter. **a**, Encoding accuracy from different LMs (averaged across voxels). We hypothesized that an LM with better performance in predicting brain responses to visual stimuli would provide more effective intermediate representations for translating visual semantic information in the brain into text. Accordingly, we performed an encoding analysis using features from each of the 42 LMs and found that the DeBERTa-large model consistently exhibited the best performance. Therefore, we used it in the main analysis. **b**, Feature correlations between target decoded features and features of descriptions generated with different MLM models. Different MLM models were used in the text generation analysis (the DeBERTa-large model was used for feature decoding and evaluation). All models exhibited reliable performance in generating descriptions aligned with target decoded features. We decided to use the RoBERTa-large model, as it exhibited the best performance. **c**, The number of tokens in generated descriptions through optimization. **d**, **e**, The number of tokens in generated descriptions (**d**) and the example descriptions (**e**) obtained with varying strength of length penalty. We evaluated the similarity between target brain-decoded features and features of candidate descriptions based on Pearson correlation coefficients with an exponential penalty to the length of candidate descriptions (see Methods for details). To determine the length penalty parameter, we performed a validation analysis with a randomly selected subset of 50 samples from the training perception data using six possible penalty parameters (0, 0.05, 0.1, 0.15, 0.2, and 0.25). While the length of generated descriptions varied depending on the length penalty, accurate descriptions of viewed content were consistently generated. We decided to use ***α*** = 0.1, as the mean length of generated descriptions with this parameter (mean = 15.56) was comparable with that of reference captions in the training perception data (mean = 15.94). In (**b−e**), the text generation analysis was performed for 50 videos randomly selected from the training perception data (DeBERTa-large; decoded from whole brain activity). Shades in (**a**, **b**) indicate 95% C.I. across subjects (*n* = 6). Error bars in (**a**, **b**) and shades in (**c**) indicate 95% C.I. across samples (*n* = 50).

**Extended Data Fig. 3.**
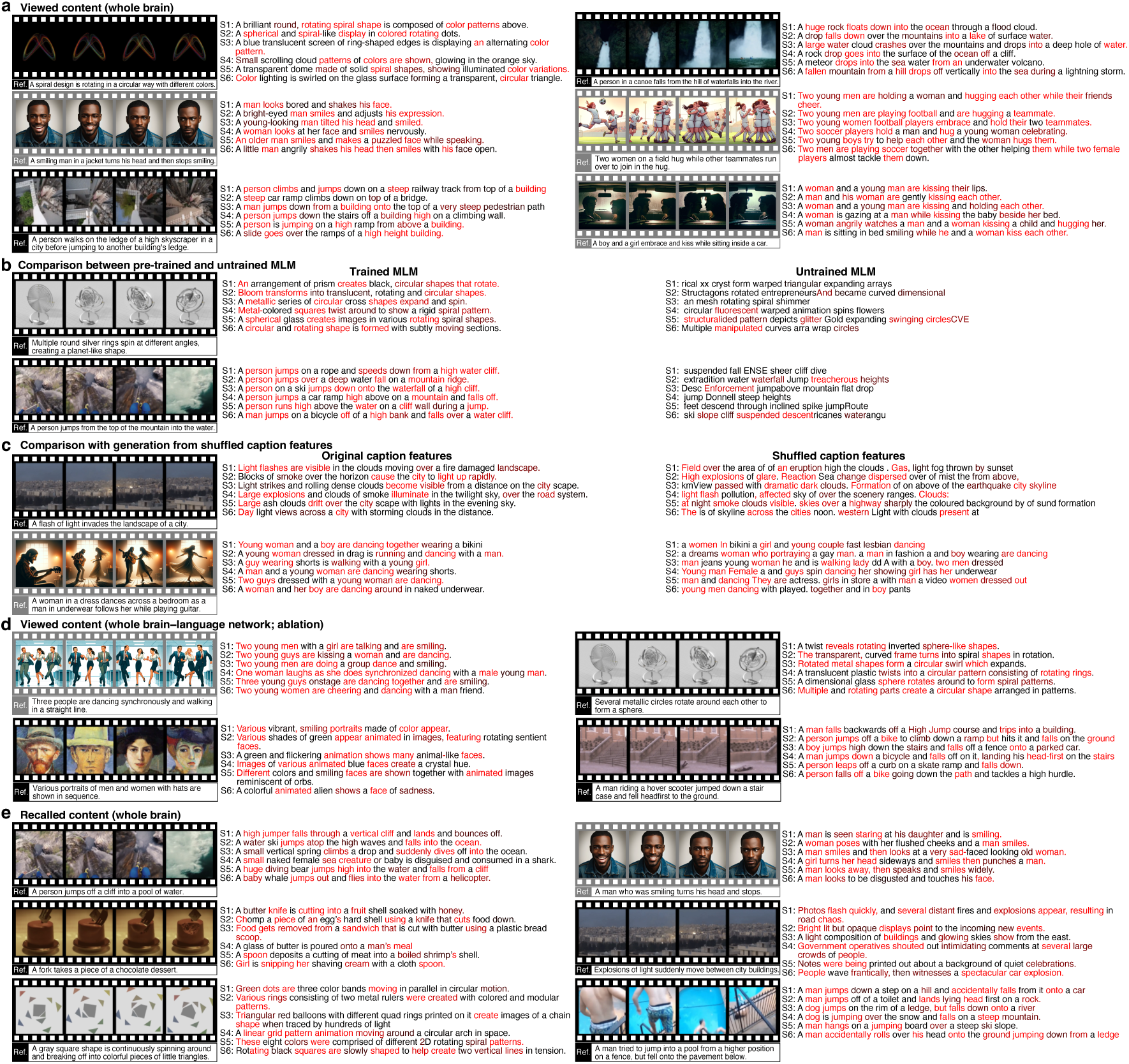
Examples of generated descriptions. **a**, Descriptions of viewed content generated from whole-brain activity. **b**, Descriptions of viewed content generated with the support of either the pre-trained or untrained MLM model. **c**, Descriptions of viewed content generated with features predicted by decoders trained based on original captions or those based on captions whose word order was randomly shuffled within each caption. Notably, the descriptions generated for the shuffled condition contained words representing individual components in viewed videos, indicating that semantic features of shuffled captions still convey the word-level semantic information. However, these descriptions lacked the coherence to accurately describe the relations among individual components. **d**, Descriptions of viewed content generated from the activity of the whole brain except for the language network. **e**, Descriptions of recalled content generated from whole-brain activity. Conventions are the same as for Fig. 2a.

**Extended Data Fig. 4.**
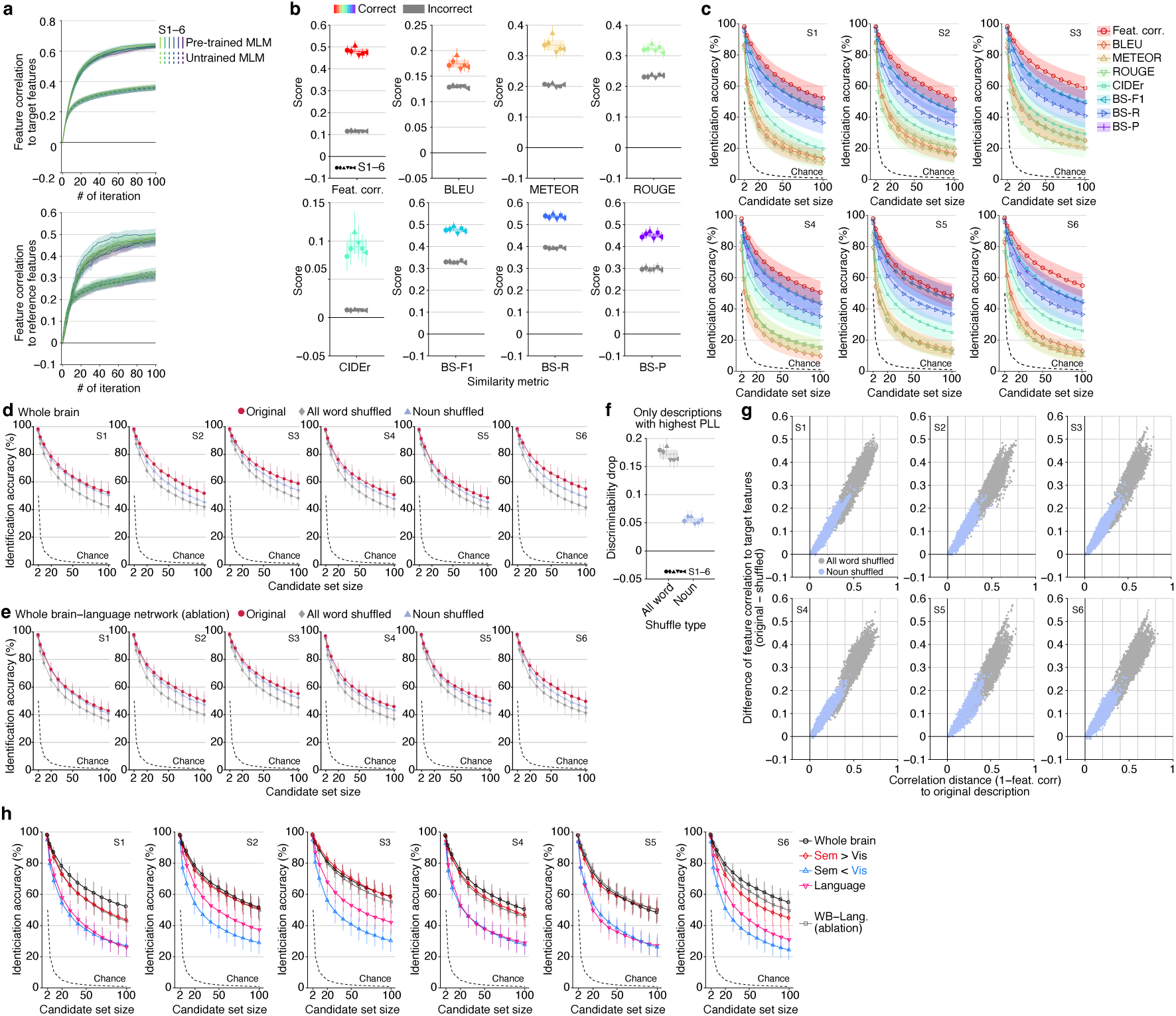
Text generation performance of viewed content for individual subjects. **a**, Feature correlations between semantic features of generated descriptions and those decoded from the brain, as well as those computed from correct references. The generated descriptions supported by the pre-trained MLM model outperformed those supported by the untrained MLM model for all subjects in both similarity measures (Wilcoxon signed-rank test, one-tailed, *P* < 0.01, FDR corrected across subjects; 100 iterations). **b**, Raw scores of the similarity between generated descriptions and captions for correct and incorrect references. **c**, Video identification accuracy. **d**, **e**, Effects of word-order shuffling on video identification accuracy applied to descriptions generated from whole-brain activity (**d**) and the activity of the whole brain except the language network (**e**). **f**, Effects of word-order shuffling on discriminability using minimally disrupted shuffled descriptions. For each generated description, reductions in discriminability were evaluated by comparing it to the shuffled variants that had the highest pseudo-log-likelihood (PLL) scores, as determined by MLM scoring^42,43^. These descriptions were selected from a pool of up to 1,000 shuffled variants created under the all-word or noun-only shuffling conditions. **g**, Scatterplot of the correlation distances (one minus feature correlation) between the original and shuffled descriptions against the difference in feature correlations to target features between the original and shuffled descriptions. **h**, Video identification accuracy obtained from different brain areas. Shades in (**a**, **c**), and error bars in (**b**, **d**, **f**, **h**) indicate 95% C.I. across samples (*n* = 72). Shades in (**b**, **f**) indicate 95% C.I. across subjects (*n* = 6).

**Extended Data Fig. 5.**
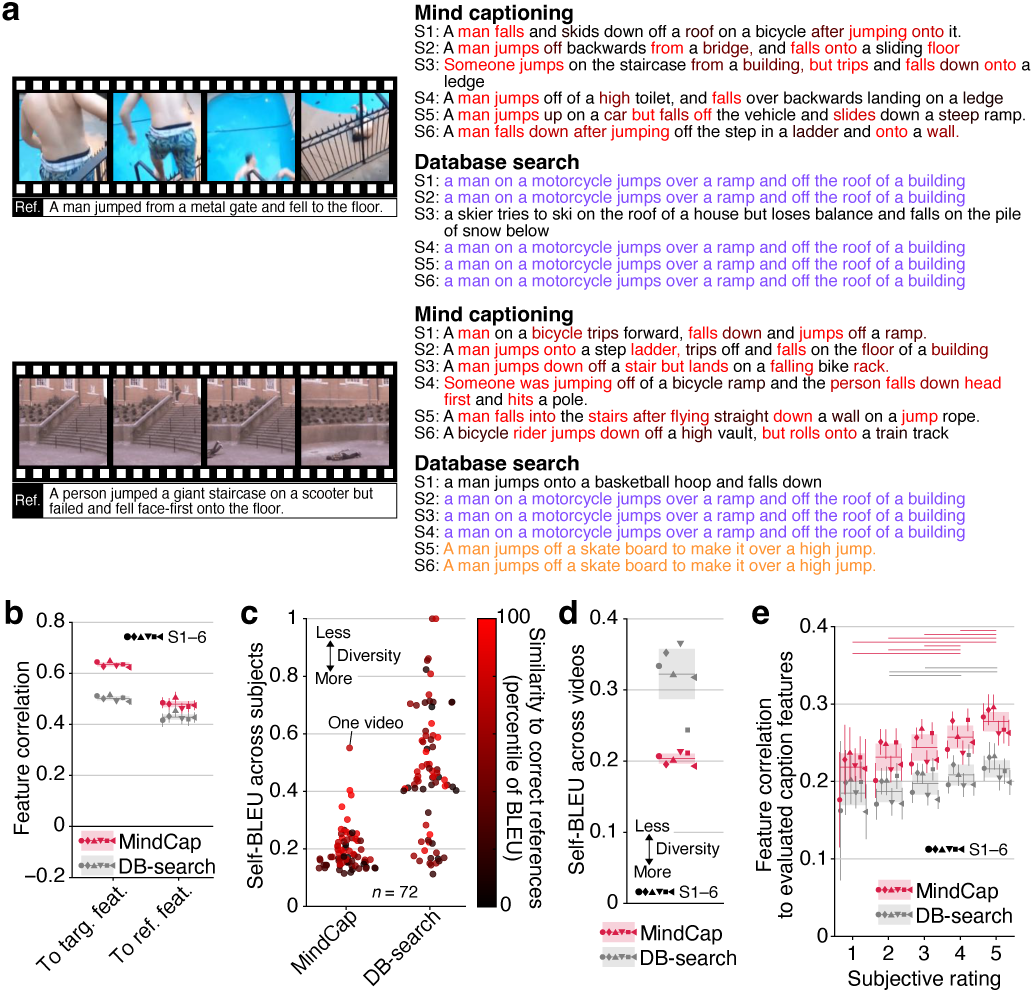
Comparison with database-search method. Text generation performance for viewed content was compared between our method and the database (DB)-search method^17^. **a**, DB-search produced moderately accurate descriptions, but it often produced identical ones for different individuals and videos. **b**, Our method outperformed DB-search, showing higher feature correlations with both target and reference features (Wilcoxon signed-rank test, one-tailed, *P* < 0.01, FDR corrected across subjects). **c**, **d**, Diversity (self-BLEU) across subjects (**c**) and videos (**d**). Our method exhibited greater diversity than DB-search both across subjects and videos (Wilcoxon signed-rank test, one-tailed, *P* < 0.05). **e**, Feature correlations at each subjective rating. Both methods showed higher scores for captions with higher ratings (lines above, *P* < 0.01, FDR corrected across pairs, six subjects pooled). Our method showed a higher correlation between ratings and feature correlations than DB-search (mean *r* = 0.184 and 0.134 for mind captioning and DB-search, respectively; six subjects averaged; *P* < 0.01, six subjects pooled), demonstrating higher flexibility in generating descriptions better aligned with subjective perceptions (ANOVA, interaction between methods and ratings, *P* < 0.01; six subjects pooled). Shades and error bars indicate 95% C.I. across subjects and samples (*n* = 6 and 72, respectively).

**Extended Data Fig. 6.**
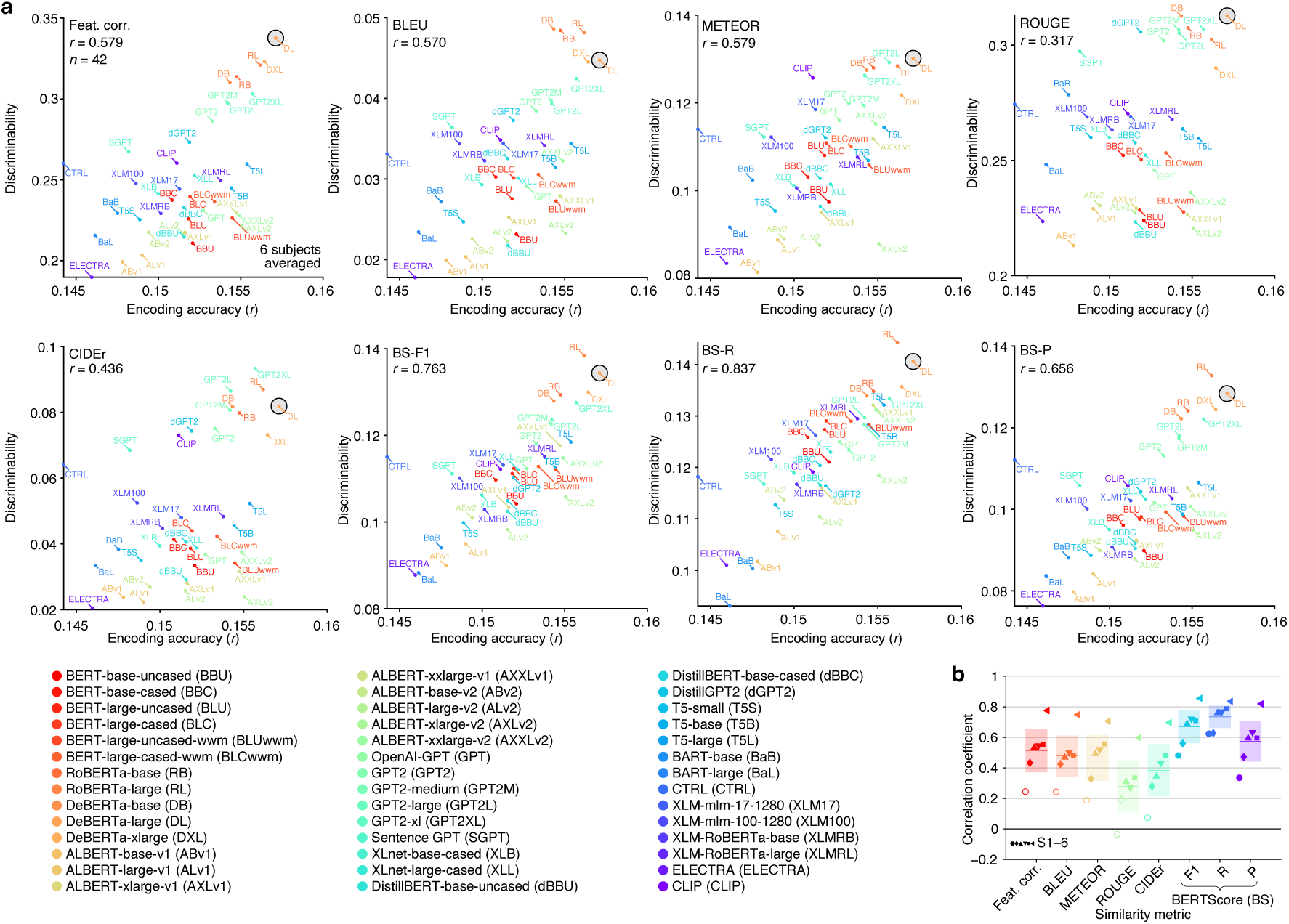
Relations between encoding accuracy and text generation performance of multiple LMs. We performed decoding and text generation analyses using features from 42 LMs. We consistently used the DeBERTa-large model for feature correlation evaluation and the RoBERTa-large model to support text generation, respectively. Encoding analysis was performed by training models on the training perception data and testing them on the test perception data. **a**, Scatterplot of encoding accuracy against discriminability (*n* = 42; dark circle, the model used in the main analysis). For each model, discriminability scores were averaged across samples, and encoding accuracy was averaged across voxels in the whole brain. The discriminability scores were positive for all models, indicating the robustness of our method to the choice of LMs. **b**, Correlation coefficients between discriminability and encoding accuracy for all models. We found significantly positive correlations for most metrics and subjects, except S1 (filled markers, *t*-test, one-tailed, *P* < 0.01, FDR corrected across metrics and subjects), suggesting that LMs with features more aligned with brain representations could help generate descriptions that accurately capture the viewed video content. Shades and error bars indicate 95% C.I. across samples (*n* = 72) and subjects (*n* = 6), respectively.

**Extended Data Fig. 7.**
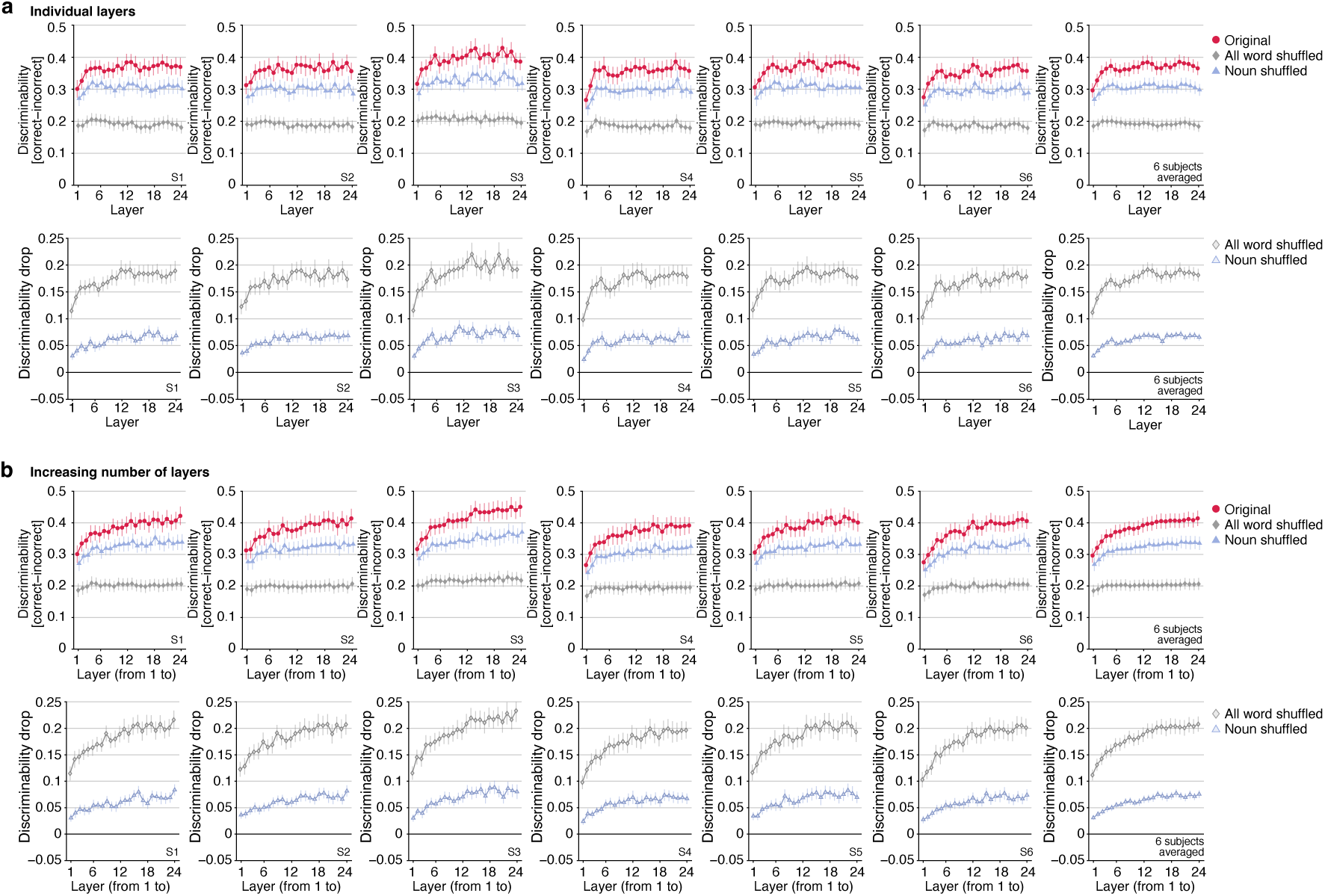
Effects of shuffling on discriminability with varying numbers of layers. The text generation analysis was conducted using decoded features with varying numbers of LM layers (24 layers from DeBERTa-large; individual layers or increasing number of layers). We evaluated the impact of shuffling on the discriminability of the descriptions generated with the varying number of layers. To compare the results from different layer conditions, we consistently used the feature correlations computed using all layers of the DeBERTa-large model as a fixed metric. **a**, **b**, Discriminability and the drop in discriminability caused by shuffling for descriptions generated with individual layers (**a**) and with increasing number of layers (**b**). Overall, text generation with deeper layers tended to result in higher discriminability (mean *r* = 0.087 for (**a**); mean *r* = 0.189 for (**b**); *t*-test, one-tailed, *P* < 0.05, FDR corrected across subjects). Furthermore, the drop in discriminability caused by shuffling became significantly larger when deeper layers were included in the analysis. Error bars indicate 95% C.I. across samples (*n* = 72; mean *r* = 0.180 [all] and 0.187 [noun] for (**a**); mean *r* = 0.259 [all] and 0.214 [noun] for (**b**); *t*-test, one-tailed, *P* < 0.05, FDR corrected across subjects).

**Extended Data Fig. 8.**
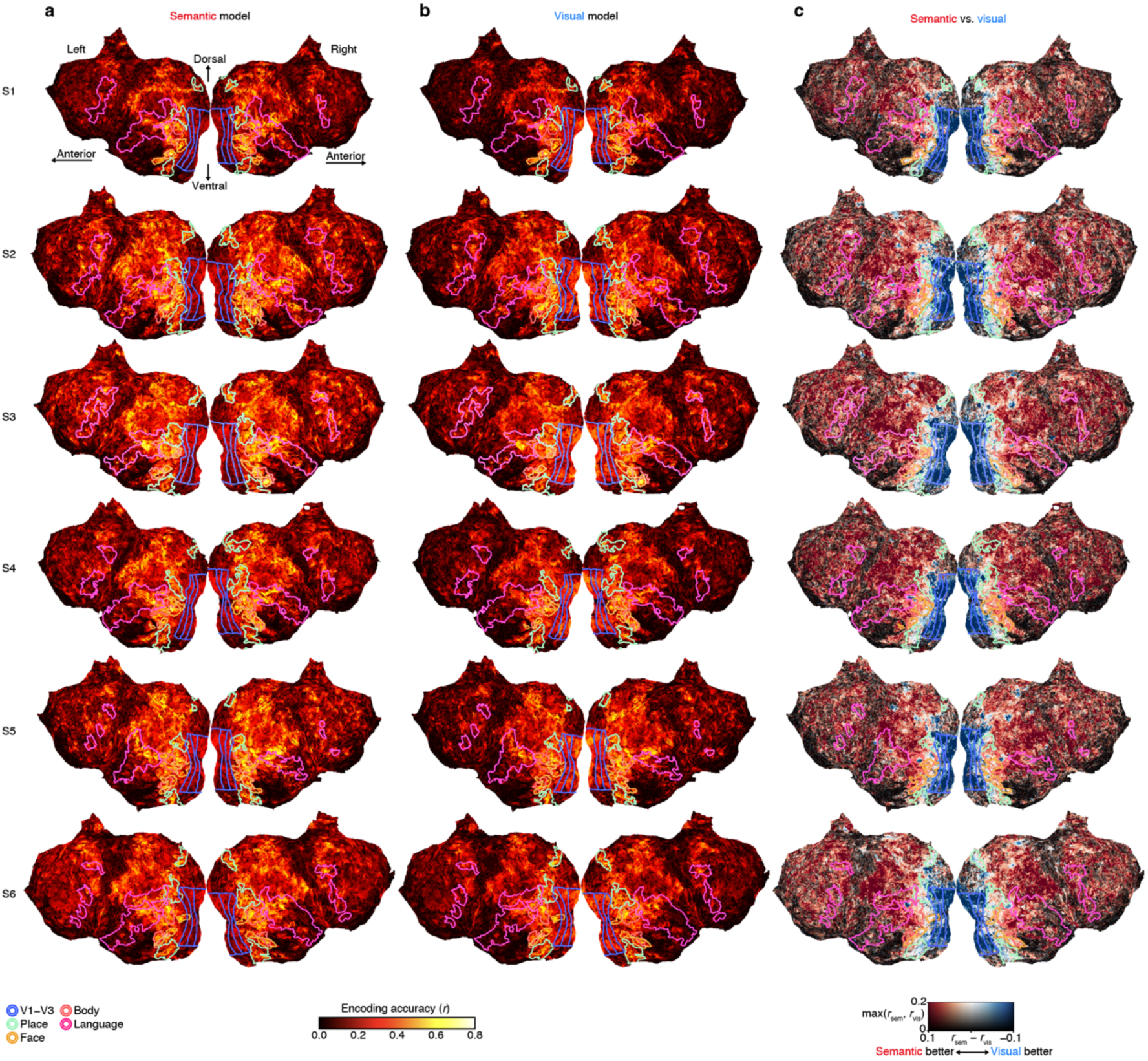
Voxel-wise encoding model performance. **a**, Encoding accuracy of the semantic encoding model. **b**, Encoding accuracy of the visual encoding model. **c**, Comparison of encoding accuracy between the semantic encoding model and visual encoding model. Across all subjects, a consistent pattern emerged in which voxels in the occipital visual areas were more accurately predicted by the visual model, whereas voxels in more anterior regions were better predicted by the semantic model. The shift in model superiority occurs at the midpoint of the category-selective regions, between their posterior and anterior halves.

**Extended Data Fig. 9.**
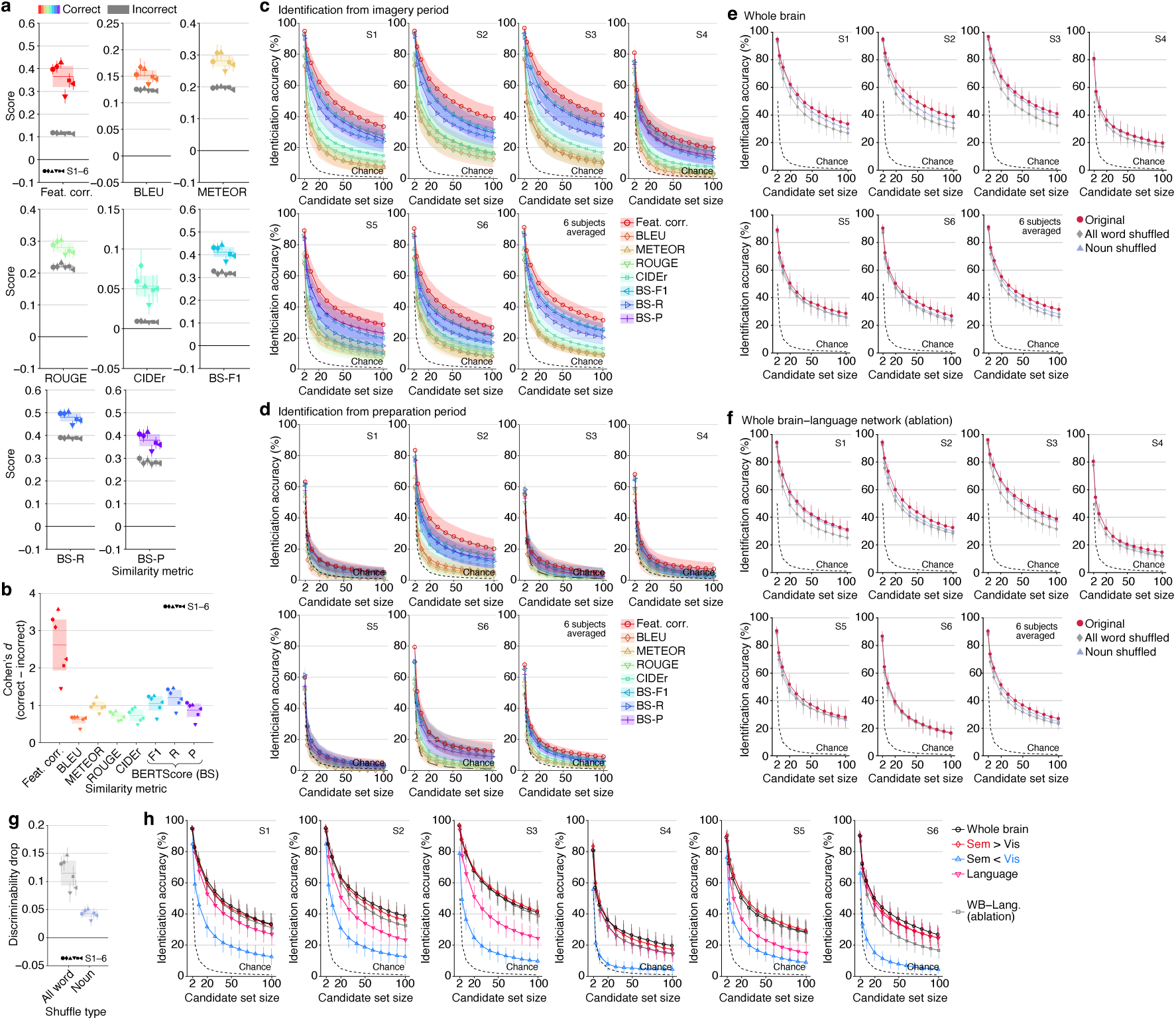
Text generation performance of recalled content for individual subjects. **a**, Raw scores for the similarity between generated descriptions and captions for correct and incorrect references based on multiple similarity metrics. **b**, Cohen’s *d* of the discriminability. The generated descriptions showed significantly high discriminability across all metrics and subjects (Wilcoxon signed-rank test, one-tailed, *P* < 0.01, FDR corrected across metrics and subjects). **c, d**, Identification accuracy of recalled videos from brain activity during the imagery period (**c**) and the preparation period (**d**). **e**, **f**, Effects of word-order shuffling on identification accuracy of recalled videos applied to descriptions generated from whole-brain activity (**e**) and the activity of the whole brain except the language network (**f**). **g**, Effects of word-order shuffling on discriminability. The word-order shuffling resulted in significant drops in discriminability even without using the language network (Wilcoxon signed-rank test, one-tailed, *P* < 0.01, FDR corrected across subjects). **h**, Identification accuracy of recalled videos from different brain areas. Shades in (**c**, **d**) and error bars in (**a**, **e**–**h**) indicate 95% C.I. across samples (*n* = 72). Shades in (**a**, **b**, **g**) indicate 95% C.I. across subjects (*n* = 6).

**Extended Data Fig. 10.**
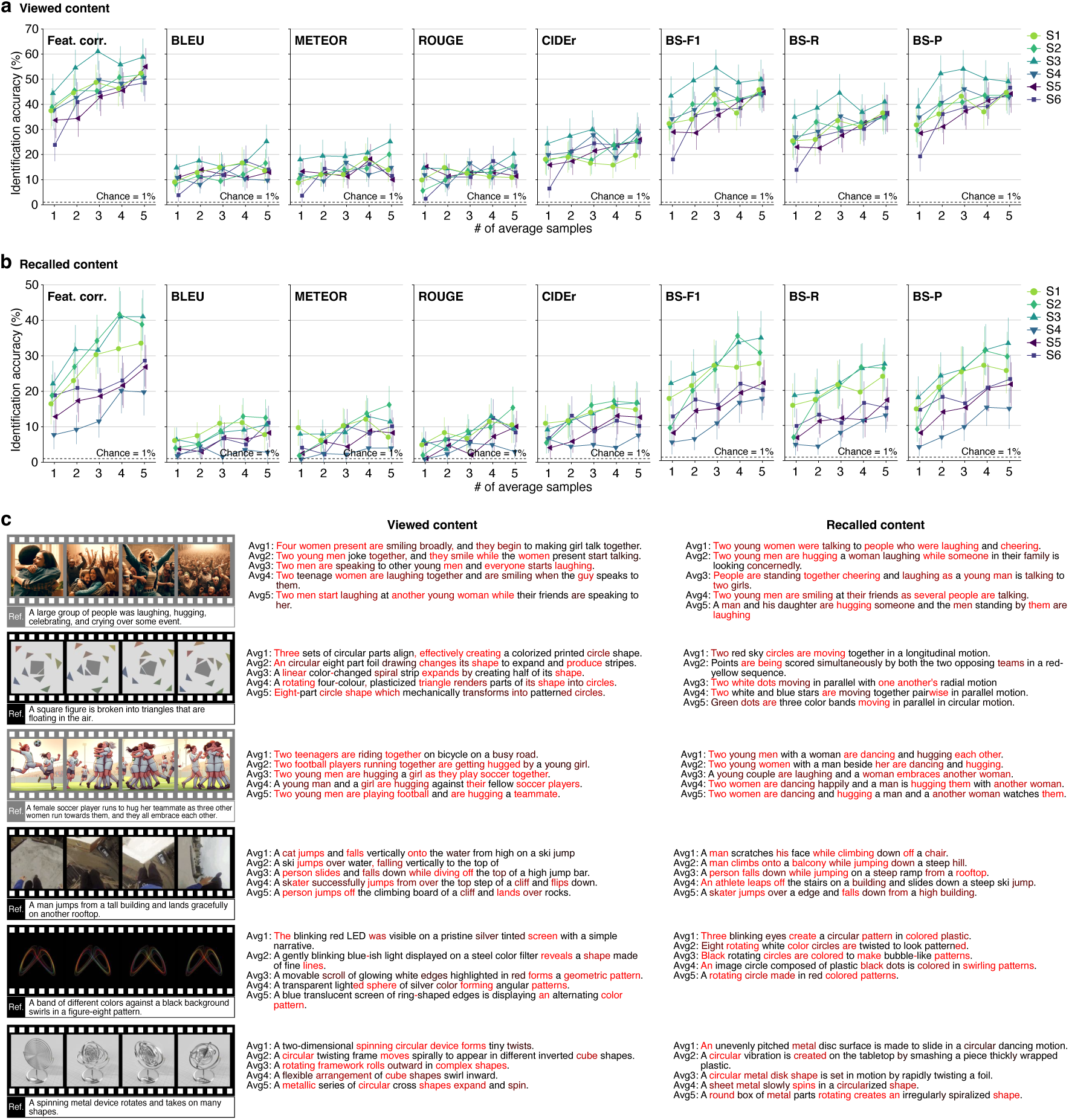
Performance of text generation with different numbers of averaged samples. The feature decoding and text generation analyses were performed with brain activity averaged across different numbers of samples. **a**, **b**, Video identification accuracy of viewed (**a**) and recalled (**b**) content with varying numbers of averaged samples. The video identification analysis was performed with 100 candidates (chance = 1%; error bars, 95% CI across samples). **c**, Descriptions generated using different numbers of averaged samples. For each video, generated descriptions from the same subjects are shown for both viewed and recalled content. The analysis revealed that the quality of the generated descriptions improved as the number of averaged samples increased, indicating that the performance was limited by noise in the fMRI signal of the test data. Meanwhile, the descriptions generated from fMRI activity in single trials reasonably captured the content of both viewed and recalled videos. These results suggest that our method is capable of generating moderately accurate descriptions of both viewed and recalled content, even when using single-trial fMRI activity.

**Supplementary Video 1. Examples of evolved descriptions of viewed content during the optimization process.** The video shows the iterative optimization process for generated descriptions of viewed content (100 optimization steps). These descriptions were generated using features from all 24 layers of the DeBERTa-large model, decoded from whole-brain activity. Conventions are the same as for Fig. 2a.

**Supplementary Video 2. Examples of generated descriptions of viewed content for all subjects.** These descriptions were generated using features from all 24 layers of the DeBERTa-large model, decoded from whole-brain activity. The text color indicates precision-based accuracy (IDF-weighted BERTScore-P).

**Supplementary Video 3. Examples of generated descriptions of recalled content for all subjects.** These descriptions were generated using features from all 24 layers of the DeBERTa-large model, decoded from whole-brain activity. The text color indicates precision-based accuracy (IDF-weighted BERTScore-P).

Supplementary Videos 1–3 are available from figshare: https://doi.org/10.6084/m9.figshare.25808179

## Notes

https://doi.org/10.18112/openneuro.ds005191.v1.0.2

https://doi.org/10.6084/m9.figshare.25808179

https://doi.org/10.5281/zenodo.15686864

### Competing Interest Statement

The authors have declared no competing interest.

### Summary of Updates

In compliance with the policy of the journal currently reviewing this work, we have reverted the preprint to the originally submitted version prior to peer review.

